# Mechanistic Regulation of Planarian Shape During Growth and Degrowth

**DOI:** 10.1101/2023.09.15.557968

**Authors:** Jason M. Ko, Waverly Reginato, Daniel Lobo

## Abstract

Adult planarians can grow when fed and degrow (shrink) when starved while maintaining their whole-body shape with correct proportions. Different planarian morphogens are expressed at the anterior-posterior poles, medio-lateral border, and midline to provide positional information signals for the specification of different tissues at the right locations. However, it is currently unknown how these signals are coordinated together during feeding or starvation and how they modulate the differential tissue growth or degrowth necessary to form correct whole-body shapes. Here we investigate the dynamics of planarian shape during growth and degrowth together with a theoretical study to evaluate the mechanisms that regulate whole-body proportions and shape. We found that the planarian body proportions scale isometrically following similar linear rates during growth and degrowth, but that fed worms are significantly wider than starved worms. By combining a descriptive model of planarian shape and size with a mechanistic model of anterior-posterior and medio-lateral signaling calibrated with a novel machine learning methodology, we demonstrate that the feedback loop between these positional information signals and the shape they control can regulate the planarian whole-body shape during growth. Furthermore, the model can predict the correct shape and size dynamics during degrowth due to an increase in apoptosis rate and pole signal during starvation. These results offer mechanistic insights into planarian shape and size dynamics and the regulation of whole-body morphologies.

## 1. Introduction

Multicellular organisms regulate the growth of their whole body across different axes to maintain appropriate shapes, sizes, and proportions, a process not completely understood (Harmansa & Lecuit, 2021; Kicheva & Briscoe, 2023; Texada et al., 2020). Planarians are ideal model organisms for studying shape regulation due to their extraordinary plasticity to grow or degrow (shrink) their adult bodies as they feed or starve, respectively (Birkholz et al., 2019; Felix et al., 2019; Lobo et al., 2012). During growth and degrowth, the planarian body dynamically changes its length at a linear rate (Almuedo-Castillo et al., 2014; Oviedo et al., 2003; Schad & Petersen, 2020) and correspondingly its body area at an exponential rate (González-Estévez, Felix, Rodríguez-Esteban, et al., 2012a; Thommen et al., 2019). Although several signaling pathways in planarians are essential to balance mitosis and apoptosis depending on nutrition intake (González-Estévez, Felix, Rodríguez-Esteban, et al., 2012b; Miller & Newmark, 2012; Pascual-Carreras et al., 2020, 2021; Peiris et al., 2012; Ziman et al., 2020), how their body proportions are dynamically regulated during growth and degrowth and which mechanism coordinates this differential growth across the axes remains poorly understood.

The whole-body rescaling in planarians is mainly due to changes in cell number rather than in cell size (Baguñà et al., 1990) and facilitated by a population of pluripotent stem cells called neoblasts that are distributed throughout the body of the worm (Scimone et al., 2014). During growth and degrowth, the differentiation of neoblasts results in a constant ratio of specific cell types (Hill & Petersen, 2015; Oviedo et al., 2003; Takeda et al., 2009). Crucially, neoblasts and their progeny are regulated by position control genes (PCGs) that form graded morphogen signals across the different axes (Bonar et al., 2022; Reddien, 2021; Witchley et al., 2013). PCGs in the Wnt pathway are necessary to correctly define the anterior-posterior (AP) axis (Gurley et al., 2008; Scimone et al., 2016; Stückemann et al., 2017; Sureda-Gómez et al., 2016), *slit* and *wnt5* are necessary to establish the medio-lateral (ML) axis (Adell et al., 2009; Cebrià et al., 2007; Gurley et al., 2010), and *bmp4* to pattern and coordinate the dorsal-ventral (DV) axis (Clark & Petersen, 2023; Molina et al., 2007; Reddien et al., 2007). However, mechanistically understanding how these signals coordinate differential growth across different axes remains a current challenge.

Mechanistic models—specifically, those based on mathematical descriptions of the underlying causes—have been proposed for explaining the planarian body patterning during homeostasis and regeneration. These dynamic models can provide mechanistic hypotheses for the formation of the planarian poles and AP patterning by reaction-diffusion (Meinhardt, 1982; Schiffmann, 2011), including the rescaling of head and tail patterns (Werner et al., 2015) as well as the location of fission planes (Herath & Lobo, 2020). Conversely, inhibitory signals diffusing from the worm AP-ML border can produce the planarian midline gradient forming from the ML axis (Meinhardt, 2004). In addition, models explaining head-trunk-tail patterning by PCGs and other morphogens can explain correct and aberrant body patterns after genetic, pharmacological, and surgical manipulations (Lobo et al., 2016; Lobo & Levin, 2015, 2017; Pietak et al., 2019), as described in formalized experimental datasets (Lobo et al., 2013). However, although these mechanistic models can predict the formation and reestablishment of key patterning signals controlling body morphology, even during changes in body size, no mechanistic model exists for explaining how these signals regulate and coordinate the dynamics of planarian whole-body shapes.

The regulation of biological shape is a complex problem due to the dynamic feedback loop between diffusible regulatory signals and the tissue locations where they act (Sharpe, 2017). In planarians, the PCG signals regulate growth and differentiation, which, in turn, determines the expression locations and diffusion domains of the PCG signals themselves. This feedback regulation involves dynamic elements at multiple scales—including cellular mechanical forces, tissue growth, and diffusion of signals—and hence requires of systems-level approaches for gaining a mechanistic understanding (Dillon et al., 2003; Germann et al., 2019; Marin-Riera et al., 2015; Mirams et al., 2013), a further challenge when considering whole organisms. Additionally, for such mechanistic models to precisely recapitulate experimental phenotypes, numerical values for all unknown parameters are needed. Thus, computational inference methods must be combined with the simulation of multiscale models in order to calibrate such models with relevant experimental data (Crocker et al., 2016; Francois & Siggia, 2010; Ko et al., 2022; Mousavi et al., 2021; Schnell et al., 2007; Uzkudun et al., 2015).

Here we experimentally study the dynamics of whole-body proportions in growing and degrowing planarians and demonstrate how the observed shapes can result from the feedback loop between pole-border signaling and the whole-body shape they regulate. We show how planarians scale isometrically at similar rates during both growth and degrowth yet maintaining different proportions. We propose and study a mechanistic regulatory model of planarian AP-ML body proportions based on the diffusion of pole and border signals regulating cell growth and the resultant body shape. Using a novel machine learning approach combining descriptive and mechanistic multi-level simulations, we successfully calibrated the proposed model to recapitulate the whole-body shape dynamics during planarian growth. Then, we employed the model to study how this regulatory mechanism can also explain planarian degrowth dynamics. We show how an increase in apoptosis and pole signaling can transition the growth dynamics into the observed degrowth whole-body shape dynamics during starvation. More generally, this study demonstrates how the complex feedback between diffusible signals controlling growth and the subsequent emergent tissue spatial dynamics can regulate the coordination and scaling of multicellular shapes.

## 2. Results

### 2.1. Planarian body proportions during growth and degrowth

Planarians increase their body size when fed (growth) and shrink when starved (degrowth). To understand the dynamics of their shape and body proportions, we imaged for nine weeks two isolated planarian populations that were either fed or starved, respectively. Figure 1 shows a set of representative images of growing (Fig. 1A) and degrowing (Fig. 1B) worms. Next, we analyzed the images with an in-house automatic algorithm to compute the width and height of each worm. The results show that the length and width dynamics each followed a similar linear trend during feeding or starvation of ±0.2 mm/week for length and ±0.04 mm/week for width (Fig. 1C-D). Analyzing the body proportion dynamics (Fig. 1E) revealed that the planarian body scaled isometrically, with allometric coefficients of 0.91 during growth and 0.98 during degrowth (the width being 16% of the length). This linear rate was not significantly different during feeding and starvation (p=0.624, one-way ANCOVA). However, the body proportions during feeding and starvation were significantly different (p<0.0001, one-way ANCOVA), with fed worms being 0.2 mm wider than starved worms.

**Figure 1.**
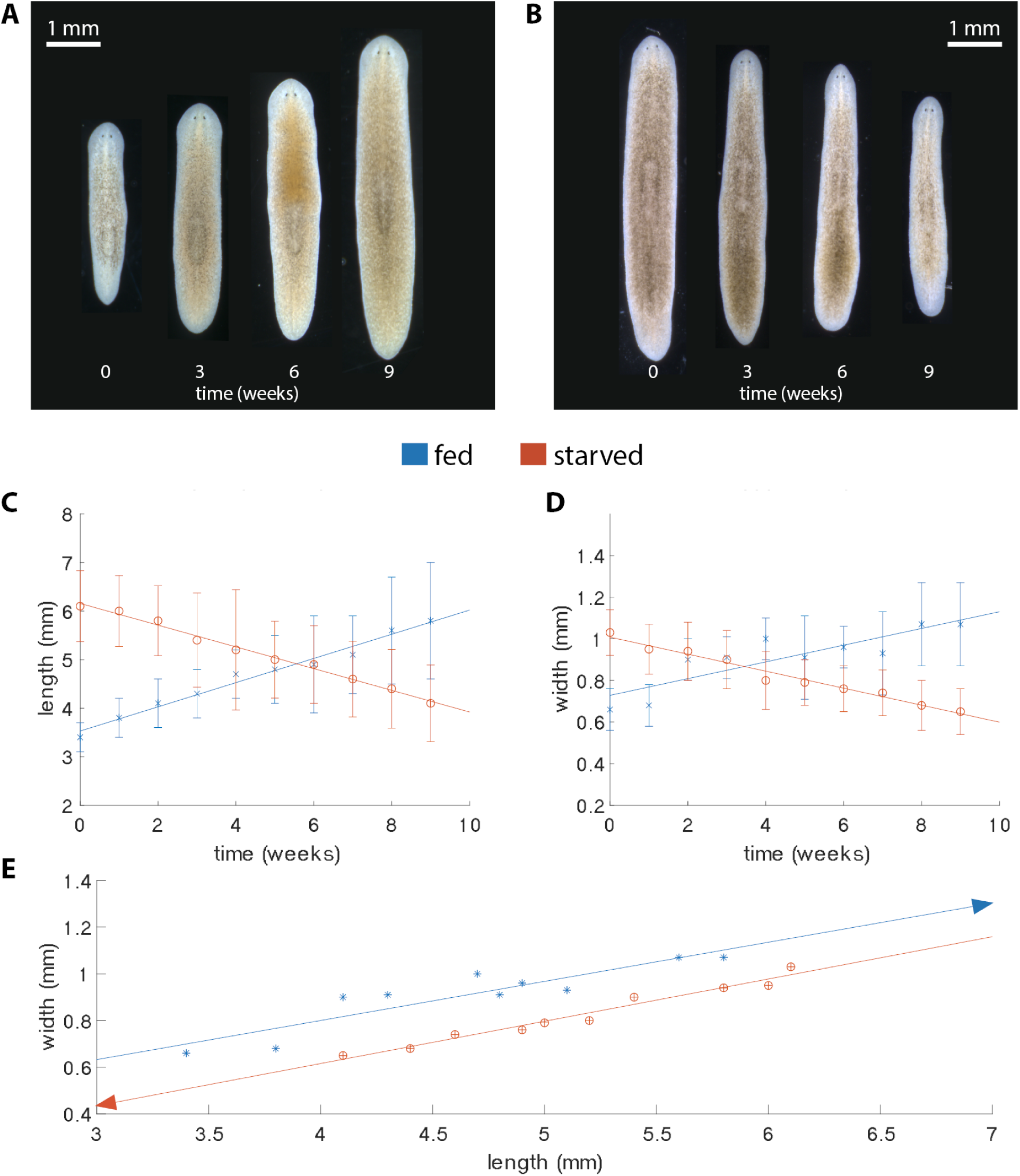
Width, length, and proportion dynamics of the planarian *Schmidtea mediterranea* during feeding and starvation. Representative images during growth when fed (A) and degrowth when starved (B). Average width (C), length (D), and proportions (E) over time during growth (blue, n=20) and degrowth (orange, n>36). Error bars indicate standard deviation.

### 2.2. Descriptive model of planarian shape dynamics

The inherent variability of the planarian body shape across different worms together with the discontinuous timing of the microscopy images (as taken once per week) hinders our ability to extract mechanistic knowledge automatically from the experimental data. To standardize the planarian body shape during growth and degrowth, we developed a continuous descriptive model of the worm shape dynamics. The model abstracts the planarian body along the anterior-posterior and medio-lateral axes as an obround geometrical shape (also called stadium), which is composed of a rectangle with two semicircular end caps (Fig. 2A). This standardized shape is specified with two parameters: the width *w* and length *l*. Then, the corresponding obround is defined as the set of all points within some radius 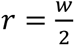 of a line segment of length *a* = *l* - 2*r*. As planarians grow and degrow, continuous obround functions can be defined using the width and length linear regression models obtained from the experimental data (Fig. 1C-D). In this way, we defined continuous standardized descriptions for the dynamic planarian shapes observed during both growth (Fig. 2B) and degrowth (Fig. 2C).

**Figure 2.**
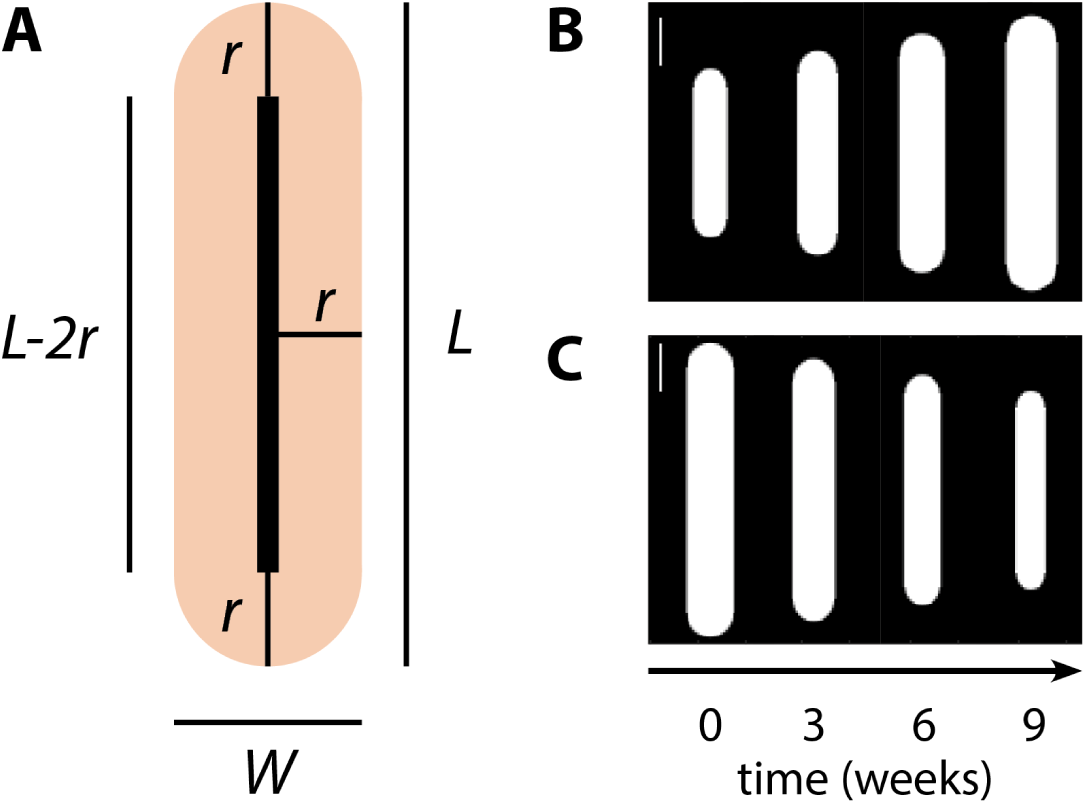
A descriptive model standardizes the planarian body shape during growth and degrowth. **A**. An obround shape with width *w* and length *l* abstracts the whole-body shape of *Schmidtea mediterranea*. **B-C**. Continuous obround functions are defined from the planarian proportion dynamics obtained experimentally during growth and degrowth, respectively. The descriptive model can thus generate planarian shapes for growth or degrowth at any timepoint (the same timepoints as in Fig. 1A-B, respectively, are shown). Scale bars: 1mm.

### 2.3. Mechanistic model of planarian shape dynamics

The descriptive model represents a standardized continuous representation of the planarian shape dynamics during growth and degrowth. Next, we sought to develop a mechanistic model that can explain the regulatory interactions controlling the dynamics of planarian shape. We hypothesized that this regulation involves the feedback between morphogen signals, controlling cell growth and division, and the existing whole-body tissue, providing a domain through which such morphogens can diffuse. To test this hypothesis, we developed a mathematical model based on partial differential equations incorporating both planarian whole-body shape and diffusible morphogen signals (Fig. 3A). We modeled the cells in the planarian body as a continuous tissue with variable cell density held together by cell-cell adhesion and through which morphogens can diffuse (Ko & Lobo, 2019). The model includes morphogens expressed at the planarian anterior-posterior (AP) pole organizers (similar to genes in the planarian Wnt pathway) and at the planarian border organizer (similar to planarian *Wnt5*). In addition, all cells can express a morphogen that function as a growth signal, inducing tissue growth. The four morphogens regulate each other in a feedforward pathway: the anterior and posterior poles (A and P in the diagram) inhibit the border morphogen (B), which in turn inhibits the growth morphogen (G). This simple regulatory pathway can establish the planarian AP and ML positional information gradients and produce a higher growth morphogen expression along the planarian midline and low elsewhere, inducing the characteristic planarian elongated whole-body shape. Crucially, the whole-body shape controlled by the morphogens feeds back into the patterns formed by the morphogens themselves, since their diffusion domain and the location of the organizers directly depend on the worm shape and size. This feedback loop could regulate the spatial dynamics of the planarian shape during growth and degrowth.

**Figure 3:**
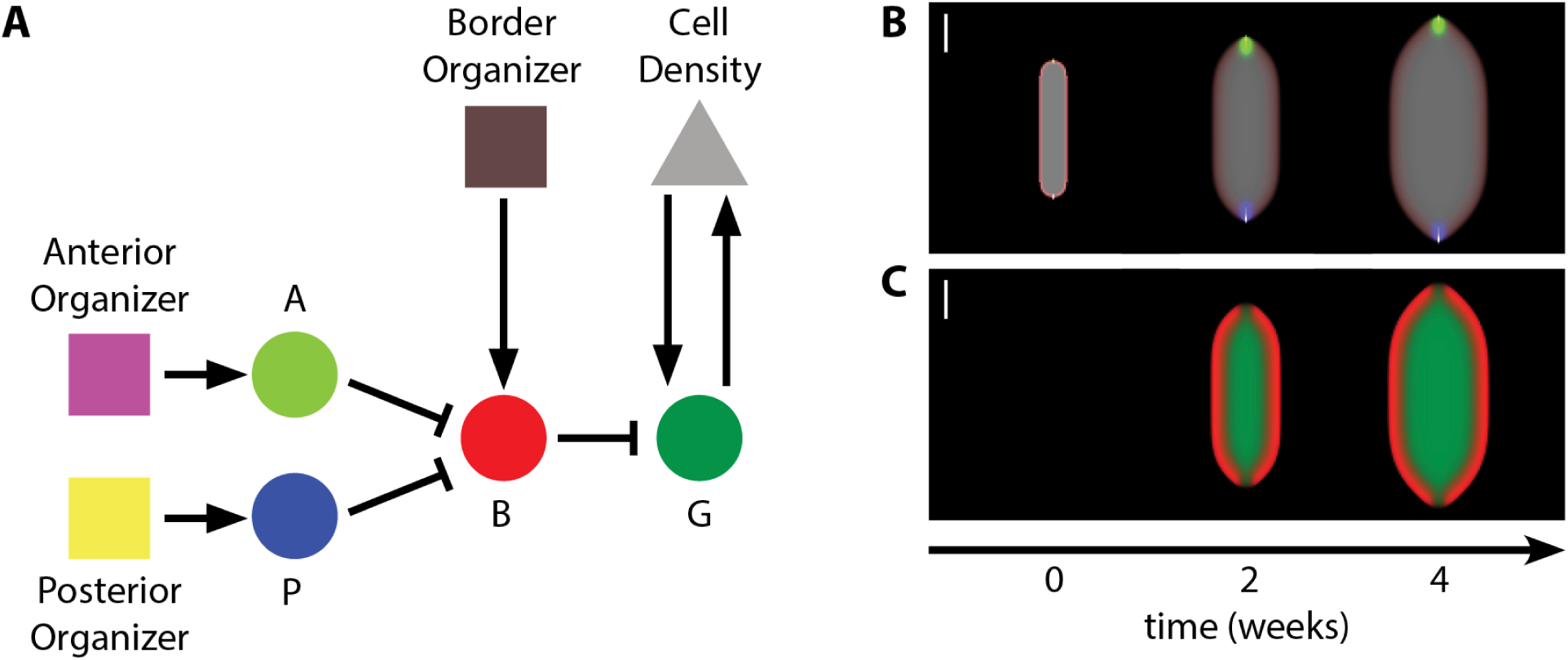
Proposed continuous mechanistic model of planarian whole-body shape. **A.** Diagram of the organizers (square), morphogens (circle), and cells (triangle) and their regulatory interactions. Organizers at the anterior (purple), posterior (yellow), and border (brown) locations express morphogens forming anterior (lime), posterior (blue), and border (red) gradients, respectively. Cells (grey) can express a growth morphogen (green) that induces tissue growth. Morphogen regulation follows a feedforward pathway, where the anterior and posterior gradients inhibit the medio-lateral border morphogen, which in turn inhibits the growth morphogen. **B-C.** Time course simulation of the mechanistic model demonstrates how the feedback loop between the tissue shape and organizers (panel B) defining the domain of the morphogens (panel C) can produce both the anterior-posterior and medio-lateral positional information gradients and the necessary differential cell growth signals resulting in an elongated whole-body shape similar to a planarian worm. Colors in (B) and (C) correspond to the levels of the respective model products in (A). The simulation domain is initialized with all products at zero concentration except cell density, which is high at the domain center with a shape and size similar to the descriptive model of planarian growth. Scale bar: 1mm. See equations in methods, parameters in Supplementary Table 1, and simulation in Supplementary Video 1.

We simulated the model to test its capacity to produce anterior-posterior and medio-lateral gradients as well as the elongated growing shapes similar to planarian worms. The initial state for cell density was generated based on the shape of the descriptive model of planarian growth. The parameters for the model were obtained from the literature or manually estimated (Supplementary Table 1). Figure 3B-C shows snapshots of the model time-course simulation, demonstrating its capacity to produce growing elongating body shapes as well as anterior-posterior and medio-lateral gradients (see also Supplementary Video 1). The expression pattern of the growth morphogen is higher at the midline and decreases towards the edges as a consequence of the inhibition by the border morphogen. The anterior and posterior morphogen gradients inhibit the expression of the border morphogen, resulting in higher cell growth at the poles and hence an elongated shape. However, the resultant simulated whole-body shape deviates substantially from the descriptive model (Fig. 2B). Manually finding the parameters that can precisely recapitulate the observed planarian shape dynamics is extremely difficult due to the multiple levels of regulation and feedback loops in the model. Instead, to optimize the parameters, we developed a machine learning approach combining the descriptive and mechanistic models.

### 2.4. Optimization of model parameters with a novel machine learning approach

To automate the discovery of model parameters that can recapitulate the observed planarian whole-body shape dynamics, we developed a novel machine learning approach. The goal of the method is to find a parameter set for the mechanistic model that, when simulated, recapitulates the shape and size dynamics during nine weeks of growth as encoded by the descriptive model and hence the experimentally observed planarian whole-body dynamics. The machine learning approach is based on evolutionary computation, an optimization algorithm that mimics biological evolution (Miikkulainen & Forrest, 2021). A random initial population of model parameter sets evolves via the application of mutation, crossover, and selection operators (see methods for details). To quantify the ability of a parameter set to precisely recapitulate planarian growth dynamics, we first define the shape error as the difference between cell density in a simulation of the mechanistic model and the shape defined by the descriptive model at the same timepoint (Fig. 4A). Each parameter set in the evolving population is scored (fitness) as the simulation time before exceeding a specified shape error threshold (Fig. 4B). In this way, simulations of parameter sets with excessive errors are terminated before reaching the total nine weeks of simulation, increasing the performance of the method. The shape error threshold decreases dynamically during evolution when parameter sets are found that can complete the whole simulation period and hence stay below the current shape error threshold. Therefore, the average simulated time increases as better model parameter sets evolve but decreases when a lower shape threshold is selected after a parameter set completes the nine weeks of simulation (Fig. 4C). This dynamic shape threshold results in more stringent simulations as evolution progresses, balancing evolutionary pressure with an enhanced performance of the method.

**Figure 4.**
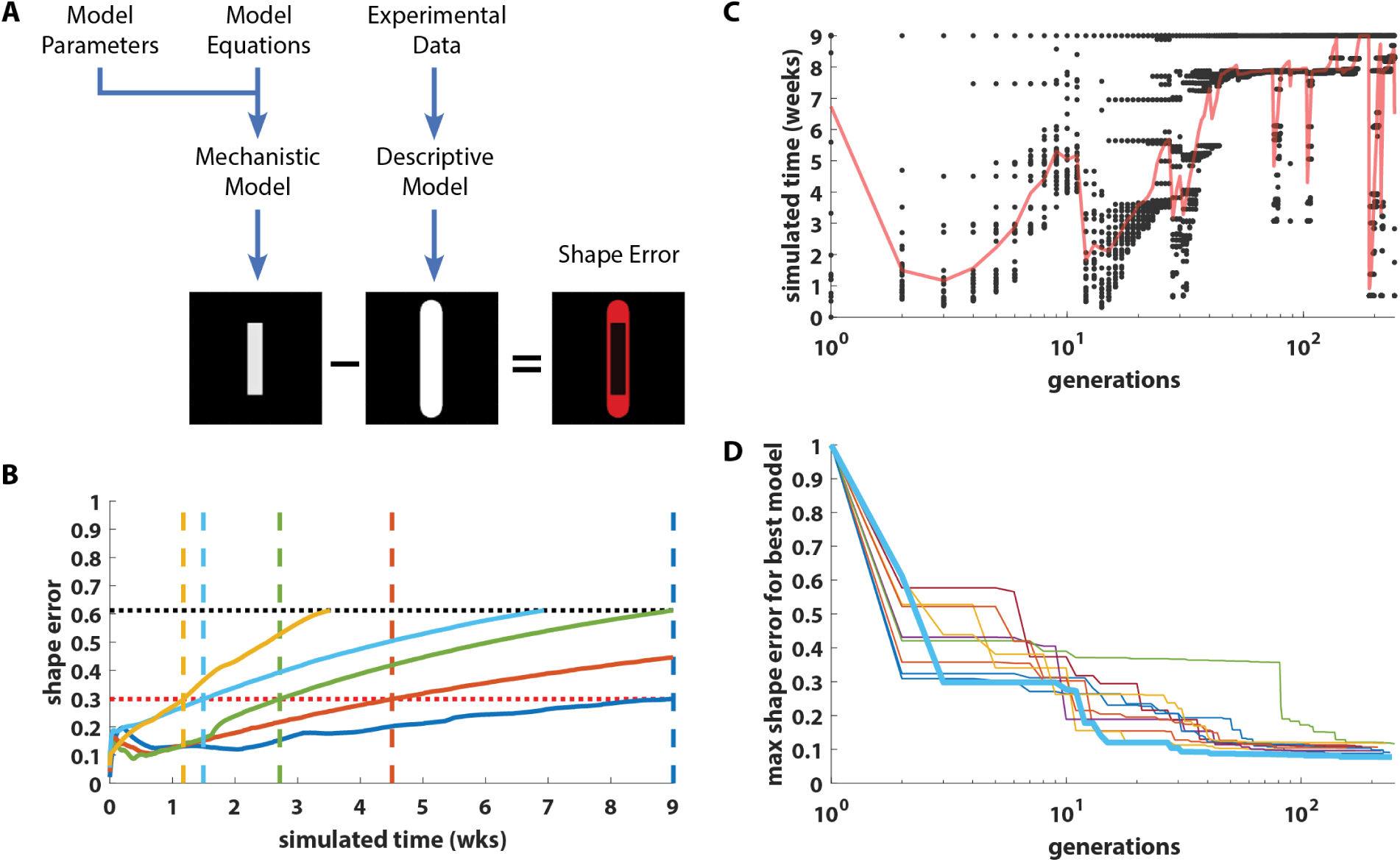
Machine learning methodology for automatically finding a set of mechanistic model parameters that can recapitulate experimental shape dynamics. A population of model parameter sets evolve until a set is found that can recapitulate the dynamics of planarian whole-body shape during growth. **A.** Shape error is computed as the difference between the simulated cell density in a mechanistic model and the descriptive model derived from experimental data at a given timepoint. **B.** Model parameter sets are simulated and scored as the time at which their shape error (solid curves) exceeds a given threshold (horizontal dotted black line). When a parameter set simulation completes the entire nine weeks (blue), its maximum shape error becomes the new shape threshold (horizontal dotted red line). **C.** Evolutionary dynamics during a run of the algorithm. Each dot represents a parameter set in the population. The average score of the evolving parameter sets (red line) improves as longer simulations complete before reaching the shape threshold. The shape threshold is updated when a parameter set can simulate for the whole nine weeks, lowering the average population score. **D.** Ten independent runs of the algorithm all produced parameter sets that can recapitulate the shape dynamics of the descriptive model of planarian growth. Best run indicated with a thick blue line.

To assess the ability of the proposed machine learning methodology to find model parameter sets that can recapitulate the planarian growth dynamics, we independently run the algorithm ten times for ten days of wall clock time each. Notice that although the mechanistic model is fully deterministic, the evolutionary algorithm includes stochasticity in the crossover and mutation operators. All ten runs converged to a parameter set with a low dynamic shape error below ∼0.1. The results demonstrate how the optimization methodology ensures that the population shape errors decrease monotonically during evolution and that all runs converged to model parameters that recapitulated the descriptive model dynamics closely (Fig. 4D).

### 2.5. Mechanistic model with discovered parameters recapitulates planarian growth dynamics

We simulated and analyzed the mechanistic model using the parameter set with the lowest error discovered by the machine learning methodology (Table 2). The time course simulation of planarian growth shows how the anterior and posterior morphogen gradients (Fig. 5A) reduce the pattern of the border morphogen at the poles (red, Fig. 5B), which in turn inhibits the growth morphogen (green, Fig. 5B) to the midline and hence away from the border except at the anterior and posterior poles. This differential growth signal results in higher growth at the poles due to the growth morphogen being closer to the border, where cell density (grey, Fig. 5A) is lower and hence causing the planarian elongated shape. Comparing the simulated mechanistic shape (Fig. 5C) with the descriptive model shape (Fig. 5D) during growth reveals a close match during the whole nine weeks of growth (Fig. 5E).

**Figure 5.**
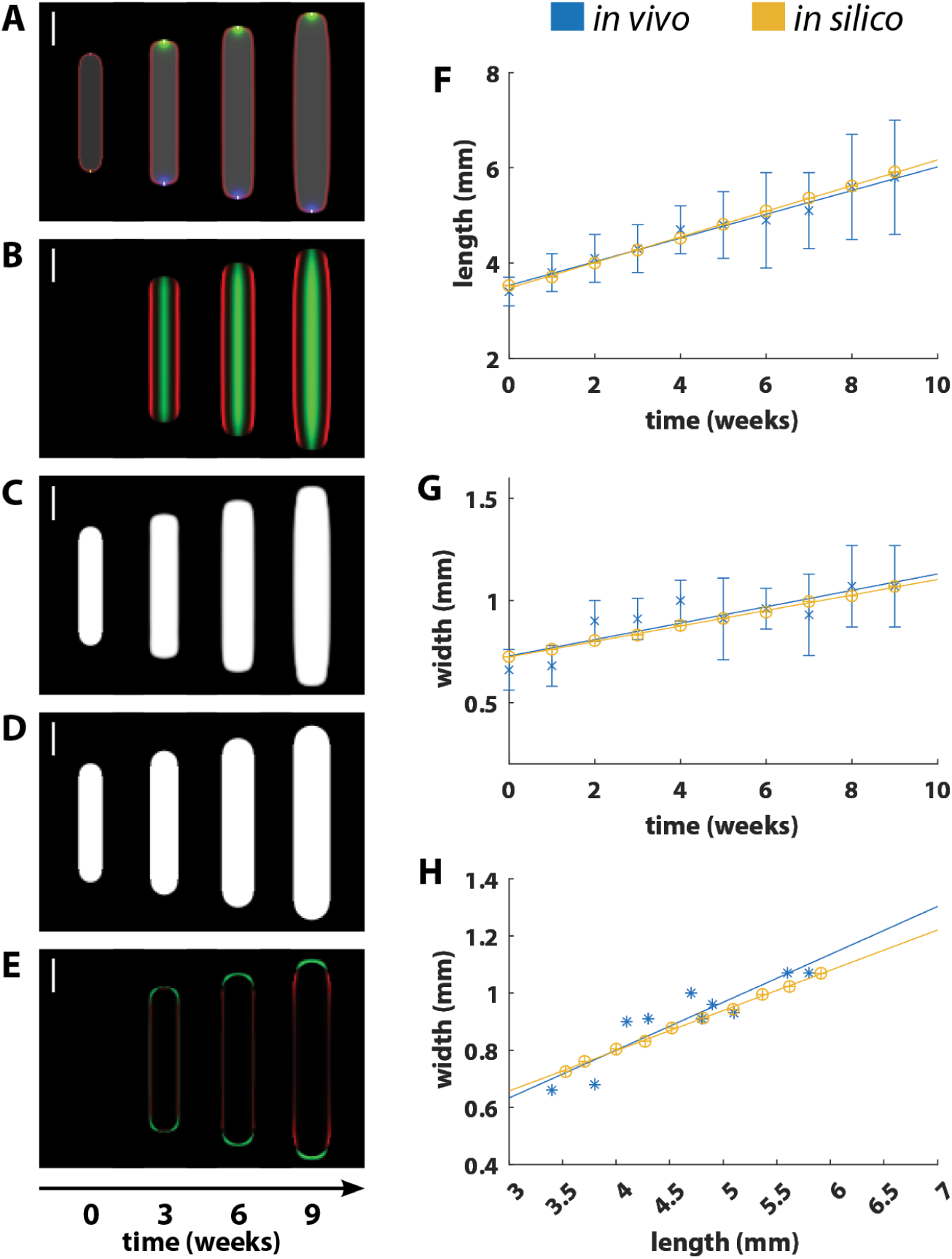
The mechanistic model with the parameter set discovered by the evolutionary algorithm recapitulates planarian growth dynamics. **A.** Anterior and posterior organizers produce pole morphogen gradients; border organizer produces border morphogen gradients. **B.** The anterior and posterior morphogens inhibit the border morphogen (red), which in turn restricts the growth morphogen (green) expression to the midline. **C.** The growth morphogen at the midline produces higher tissue growth at the border near the poles where cell density is lower, which results in an elongated growing shape. **D.** Descriptive model of planarian growth derived from experimental data. **E.** Spatially subtracting the simulations with the descriptive and mechanistic models reveals a good match between the shapes (green and red indicates excess mechanistic or descriptive shape, respectively). **F-G.** Dynamics of planarian growth *in vivo* (blue) and *in silico* (yellow) in terms of body length (F), width (G), and proportion (H), which is not significantly different. Error bars indicate standard deviation. Colors for (A-E) as in schematic in Fig. 3A. Scale bar: 1mm. See parameters in Supplementary Table 2 and simulation in Supplementary Video 2.

Next, we compared the planarian length, width, and body proportions resulting from the mechanistic model during growth with the experimental measurements of planarian worms. These results show that the mechanistic model correctly recapitulated the width and length dynamics during the nine weeks of growth experimental data (Fig. 5F-G). Furthermore, the planarian body proportion dynamics were not significantly different between the mechanistic model and experimental measurements (Fig. 5H), both in terms of rate and proportions (p=0.340 and p=0.287, respectively, one-way ANCOVA). Thus, the mechanistic model with the automatically discovered parameters is able to reproduce the planarian whole-body shape dynamics during growth.

We performed a sensitivity analysis to determine the robustness of the mechanistic model of planarian growth and the effect of each of its parameters on the resulting shape. We perturbed each parameter calibrated with the machine learning method by applying factors between 0.5 and 1.5. For each perturbation the growth simulation was repeated from the original initial state and the resultant whole-body shape analyzed after three simulated weeks (Sup. Fig. 1). The resultant whole-body shapes show how each of the individual parameters can affect different morphological aspects of the growing worm. Several parameters, such as the diffusion rate of the anterior and posterior morphogens (*D*_*AP*_) or the growth morphogen production rate constant (*b*_*G*_), can control the convexity of the shape. In addition, parameter can also affect the overall length and width of the resultant shape, such as the apoptosis rate constant (*λ*) or cell dispersion constant (*k*_*p*_). In this way, the mechanistic model is robust and can produce a variety of growing shapes depending on the values of the different biological parameters.

### 2.6. Mechanistic model with higher cell apoptosis and pole signaling recapitulates planarian degrowth dynamics

We sought to validate the calibrated model trained with the planarian experimental growth dynamics by recapitulating the observed dynamics during starvation (Fig. 1). When planarians are starved, the rate of cell apoptosis increases (Pellettieri et al., 2010) and the rate of cell mitosis decreases (González-Estévez, Felix, Rodríguez-Esteban, et al., 2012a) as compared to fed worms. Indeed, a lower rate of apoptosis *λ* decreases overall cell growth in the model, but this change alone distorts the body shape produced (Sup. Fig. 1). Hence, we hypothesized that increasing both the rate of apoptosis *λ* and the pole coefficient α (which modulates the regulation of the border morphogen by the pole morphogens) in the calibrated mechanistic model could be sufficient to cause the observed degrowth shape dynamics during starvation. To test the hypothesis, we performed a parameter scan varying both the rate of apoptosis *λ* (increasing up to 10%, at 2% increments) and the effect of the pole signaling α (increasing from −1 to 1, at 0.2 increments) for a total of 66 simulations starting with the body shape and size defined by the descriptive model for the planarian degrowth experiments (Fig. 2C).

The results of the parameter scan show how the calibrated model could produce a variety of degrowth behaviors with different body proportion dynamics (Fig. 6A). Increasing the pole coefficient α results in higher production of the border signal *B* as regulated by the pole morphogens *A* and *P* (Fig. 6B), and hence a localized lower cell growth at the poles that changes the resulting whole-body shape. The pole coefficient correlated more strongly with the resulting body length (Fig. 6C), while the rate of apoptosis influenced more strongly the body width (Fig. 6D). Furthermore, the simulations that resulted in correct body proportions also matched the body shape and size defined by the descriptive model derived from the experimental degrowth data (Fig. 6E).

**Figure 6.**
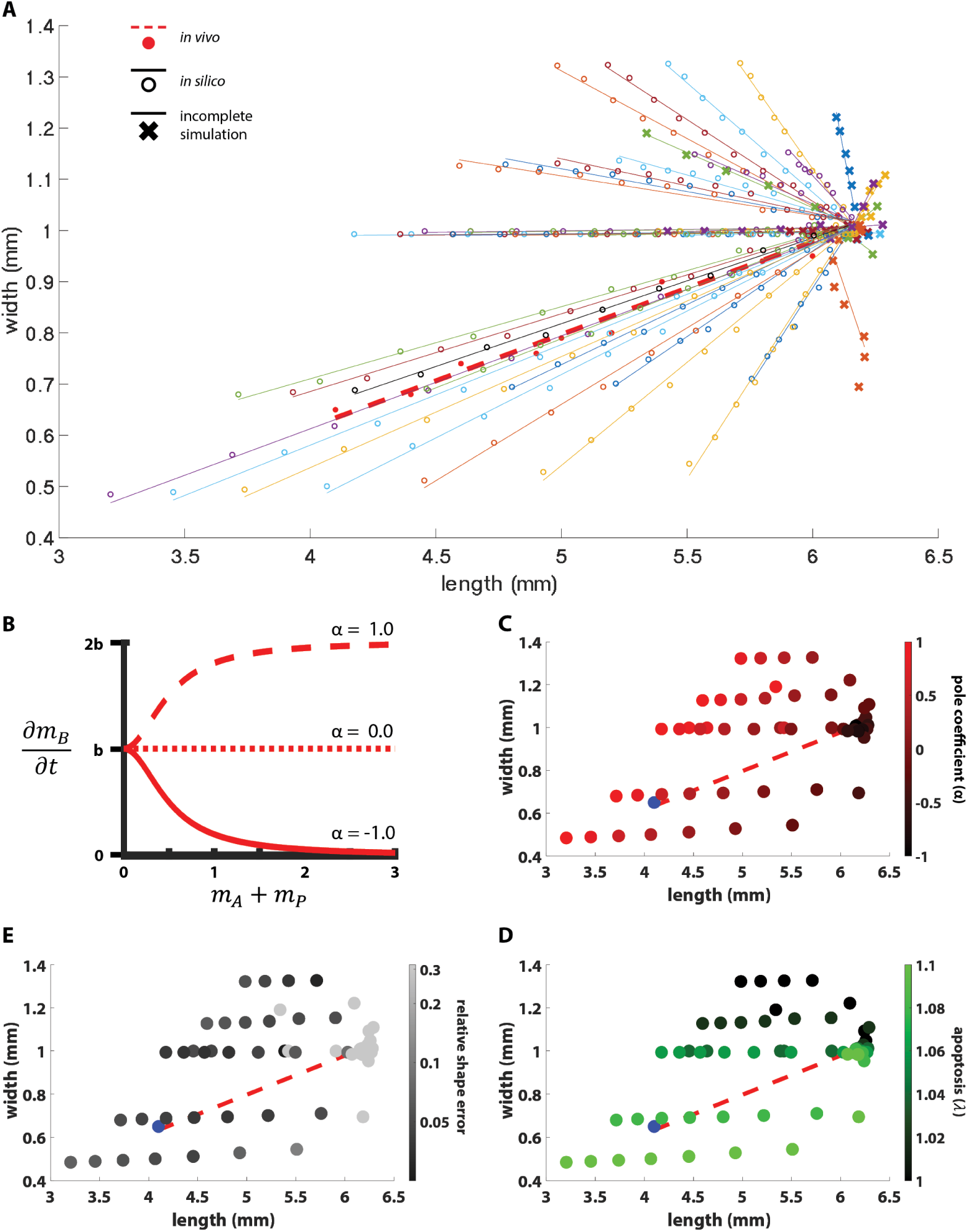
Increasing the pole coefficient and apoptosis rate in the calibrated mechanistic model of growth can produce a variety of degrowth dynamics. **A.** Dynamic shape proportions of simulations from a parameter scan (solid lines) compared to the experimental degrowth data of starved planarians (dashed red line). Best simulation (black line) matches the experimental proportion dynamics. Simulations marked with ‘x’ terminated earlier than the nine weeks of experimental data due to exceeding the simulation domain (and hence growing) or excessive stiffness. **B.** Effect of varying pole coefficient *α* in the regulation between the pole morphogens (*m*_*A*_ + *m*_*P*_) and the border morphogen *m*_*B*_. **C-E.** Varied parameter values and resulting shape error in the parameter scan. Each dot represents a simulation (final shape); dotted line represents the experimental degrowth data (final shape in blue).

Among all the simulations in the parameter scan, an 8% increase in the rate of apoptosis and an increase of the pole coefficient to 0.6 resulted in the closest match between the simulated (Fig. 6A, black line) and the experimental (dash red line) whole-body proportion dynamics during degrowth. The simulations with this model also revealed that the whole-body shapes closely matched those in the descriptive model from the experimental degrowth dynamics (Fig. 7A-E). Furthermore, the predicted dynamics with the degrowth model also matched the experimental data of planarian degrowth in terms of length (Fig. 7F) and width (Fig. 7G). In addition, the proportions of the simulated dynamic shapes (Fig. 7H) were not significantly different from those obtained experimentally in terms of both their rates (p=0.220, one-way ANCOVA) and magnitudes (p=0.036, one-way ANCOVA). Finally, we tested weather an increase in apoptosis rate together with any other parameter could induce correct degrowth dynamics. The results demonstrated that only the combination of increased apoptosis and pole coefficient could induce a decrease in both length and width and hence correct degrowth dynamics (Sup. Fig. 2).

**Figure 7.**
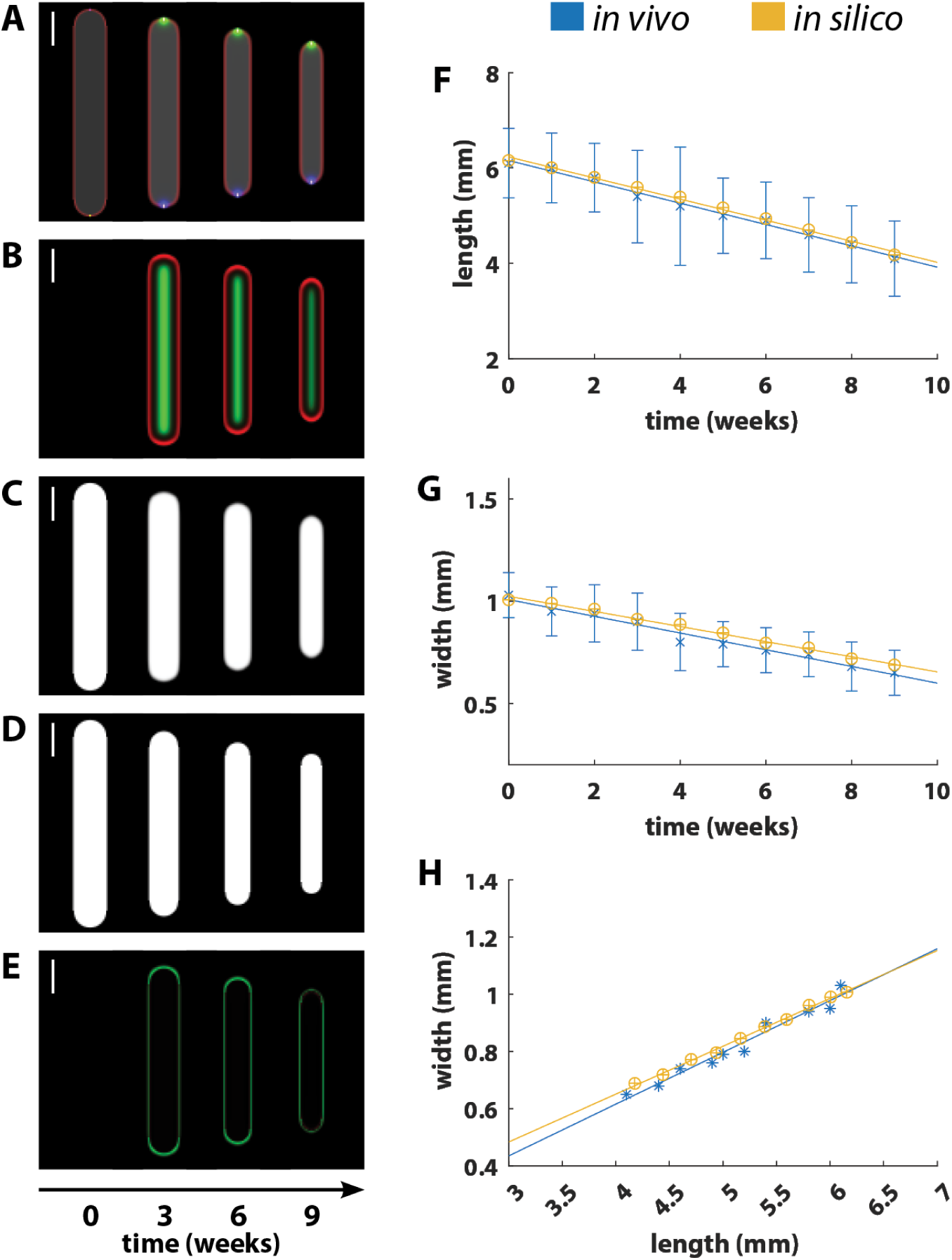
The calibrated mechanistic model with higher apoptosis rate and pole coefficient recapitulates planarian degrowth dynamics. **A.** The increase in apoptosis results in degrowth dynamics, even with similar anterior, posterior, and border organizers. **B.** The increase in pole coefficient results in higher expression of border morphogen at the poles, which locally reduces the expression of growth morphogen. **C.** The reduction of growth morphogen at the poles induces additional degrowth at these locations, which results in faster shortening than narrowing. **D.** Descriptive model of planarian degrowth derived from experimental data. **E.** Spatially subtracting the simulations with the descriptive and mechanistic models reveals a good match between the shapes (green and red indicates excess mechanistic or descriptive shape, respectively). **F-G.** Dynamics of planarian degrowth *in vivo* (blue) and *in silico* (yellow) in terms of body length (F), width (G), and proportion (H), which is not significantly different. Error bars indicate standard deviation. Colors for (A-E) as in schematic in Fig. 3A. Scale bar: 1mm. See simulation in Supplementary Video 3.

## 3. Discussion

Here, we experimentally have studied the planarian body proportion dynamics during growth and degrowth and proposed a mechanistic mathematical model of its regulation that could recapitulate the observed shapes. Employing an automated image analysis pipeline to either fed or starved planarian populations, we determined that their body proportions are significantly different— growing worms being wider than degrowing ones—yet that they scale isometrically following the same rates. Next, we sought to understand the regulatory mechanisms that could control the differential tissue growth and degrowth dynamics required to produce the observed whole-body shapes and sizes. Due to the complex feedback between shape and the expression and diffusion of signaling morphogens, we employed a multi-level mathematical approach where cell density is modeled as a dynamic continuum subjected to cell adhesion forces and that can grow and degrow in response to the concentration of morphogens. In turn, the expression locations and diffusive domains of such regulatory morphogens depend on the dynamic shape of the tissue they regulate, closing a mechanochemical feedback loop. We showed how such mechanistic model integrating the regulation of growth signals by border and AP-pole morphogens could produce obround-like shapes. To demonstrate that the model could precisely recapitulate the observed planarian shape dynamics, we developed a machine learning approach to calibrate the parameters for the mechanistic model to recapitulate the observed planarian shape dynamics. To generalize the experimental planarian shape image data taken at a resolution of once per week, we defined a continuous descriptive model that abstracted the planarian whole-body shape as an obround. Then, taking this descriptive model as input, the machine learning approach evolved a parameter set that could minimize, when simulated, the difference of whole-body shapes between the descriptive and mechanistic models. The resulting calibrated mechanistic model could precisely recapitulate the observed planarian proportions during growth.

To understand how planarians shift between regulating growth when fed to degrowth when starved, we used the calibrated mechanistic model to perform simulations increasing cell apoptosis and varying one other model parameter at a time. This parameter scan showed that an increase in cell apoptosis alone cannot explain the transition to degrowth shape dynamics during starvation. Instead, only the combination of increasing apoptosis and the pole signaling resulted in the precise degrowth dynamics that recapitulated the observed planarian proportions during starvation, suggesting that the positional control genes patterning the AP axis are actively regulated during degrowth. Thus, even though planarian proportions during growth and degrowth are significantly different, the same proposed mechanistic model serves as a combined explanation of how the planarian body shape is regulated during both growth and degrowth.

The proposed pole and border morphogens in the mechanistic model could be implemented by several planarian genes that act as axis organizers (Sureda-Gomez & Adell, 2019). In particular, diffusive Wnt ligands are expressed at the planarian anterior and posterior ends (Gurley et al., 2010; Petersen & Reddien, 2009, 2011), similar to the pole organizers proposed in the mechanistic model. The non-canonical Wnt ligand *wnt5* is expressed at the planarian AP-ML border (Cebrià et al., 2007; Gurley et al., 2010), like the border signals included in the model, and is antagonistic to medial *slit*. Finally, pathways such as EGFR and TOR control planarian cell proliferation (Fraguas et al., 2011; González-Estévez, Felix, Smith, et al., 2012; Peiris et al., 2012), and, as proposed in the model, may need to be coordinated with AP pole genes in the Wnt (and dorsoventral BMP) pathway controlling differential growth (Clark & Petersen, 2023).

Although the proposed mechanistic model can precisely recapitulate the dynamics of planarian growth and degrowth shapes and sizes, it includes simplifications to facilitate the mathematical simulations. First, the model is based on direct pathways from organizers to tissue growth. However, the equivalent planarian pathways likely involve multiple genes and signals that also participate in other processes such as cell differentiation, which is also omitted in the model. In addition, the model excludes cell migration for simplicity and instead abstracts tissue growth as the effect of both cell proliferation and migration. Indeed, the differentiation of neoblasts involve their migration towards the target tissues (Lapan & Reddien, 2011), effectively increasing cell density in those locations and hence resulting in tissue growth. Finally, the model is focused on the AP and ML axes, but future work could extend this approach to three dimensions to incorporate the important contribution of DV patterning in the regulation of planarian whole-body shapes (Gaviño & Reddien, 2011).

In conclusion, this study suggests that the planarian whole-body shape is regulated by the feedback interaction between AP-ML morphogens controlling differential growth and the emergent tissue shapes where they are expressed and diffuse. Our approach combining experimental imaging analysis, multi-level mathematical modeling, and machine learning enabled the understanding of this complex mechanochemical feedback regulating shape in planarians. Crucially, by combining a descriptive and mechanistic model, we generated an intuitive continuous definition of model error grounded in experimental data, which served as the training basis for the inference methodology. Despite using machine learning, the resultant mechanistic model is interpretable and hence could be used to shed light on plausible regulatory mechanisms controlling planarian shape during growth and to predict the process for which it can transition into degrowth. Further work will be essential to experimentally validate the predictions posed by the mechanistic model, but this work can pave the way towards a general understanding of the interrelation between morphogens and shapes in animal development.

## 4. Materials and Methods

### 4.1. Planarian culture

Asexual strain CIW4 of the planarian *Schmidtea mediterranea* were maintained in 1× Montjuic water in an incubator at 20°C in the dark as previously described (Merryman et al., 2018). Two planarian populations were kept separated, each initially composed of either small animals of ∼3mm length fed beef liver paste twice per week (N=20), or large animals of ∼6mm length starved (N=50). Planarians that fissioned in the starved population were removed (final N=36). No planarians fissioned in the growth population during the reported period.

### 4.2. Planarian measurements

Each week, three images were taken for each planarian when gliding in a straight line using a Zeiss SteREO Discovery.V20 zoom stereoscope with dark field illumination. Planarian body proportions were automatically measured for each image with a custom pipeline in MATLAB (The MathWorks, Inc.) by binary thresholding and then computing the minimum bounding rectangle of the largest connected component, so small debris and visual artifacts were discarded.

### 4.3. Mechanistic model

The mechanistic model of planarian growth and degrowth represents the planarian whole-body as a continuum of cell density *u*, where morphogen signals can be expressed, advected, and diffuse. As in (Ko & Lobo, 2019), cells are subjected to a velocity vector field, ***V***, as well as growth and apoptosis modulated by growth morphogen *m*_*G*_ as defined by function *g*(*u*, *m*_*G*_). Thus, the change in cell density over time can be expressed as

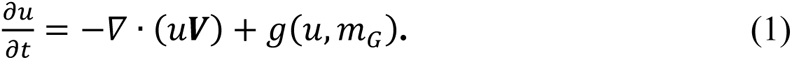

Cell velocity is comprised of two components, dispersion (***V***_***d***_) and adhesion (***V***_***a***_), such that

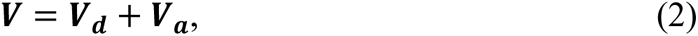

where ***V***_***d***_ is the dispersion velocity, and ***V***_***a***_ is the adhesion velocity. Diffusion is assumed to be negligible relative to the strength of dispersion and adhesion.

Dispersion is proportional to cell density, causing cells to move away from each other. As dispersion velocity is proportional to the population density,

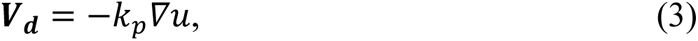

where *k*_*p*_ is the dispersion constant.

All cells within some radius R of each other are drawn together via adhesion forces, which are defined with a non-local integral term,

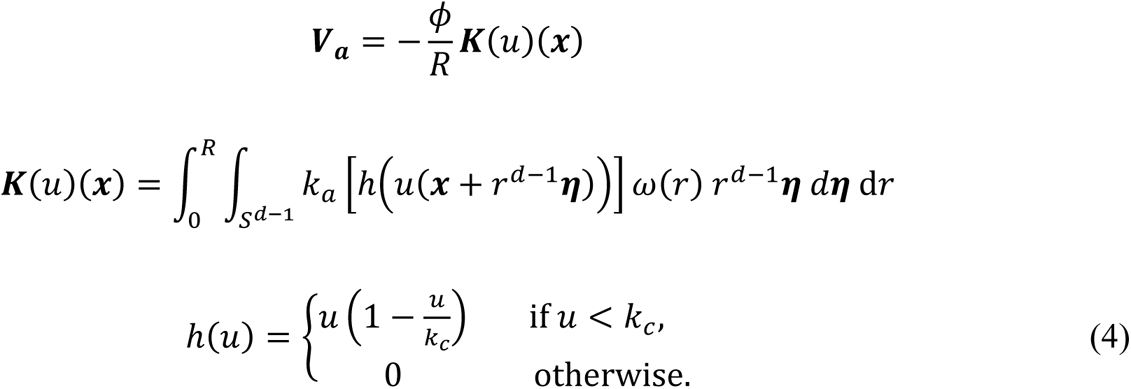

where *R* is the radius of adhesion, *ϕ* is a constant of proportionality related to viscosity, *k*_*a*_ is the adhesion strength constant, and ℎ(*u*) is a crowding capacity function which prevents cells from moving into regions of high cell density. The function *ω*(*r*) describes how the force varies with the radial distance *r*. For simplicity, we define *ω*(*r*) = 1.

The last component of the cell density equation is the function *g*(*u*, *m*_*G*_), which describes changes due to tissue growth and cell apoptosis. A growth morphogen *m*_*G*_ signals the level of tissue growth, which is subjected to the carrying capacity *k*_*c*_. Additionally, all cells experience apoptosis with rate constant *λ*. Hence,

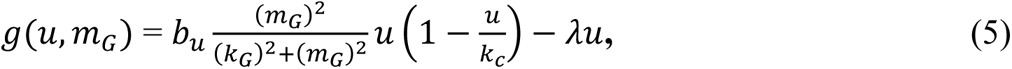

where *k*_*G*_ is the half-maximal concentration of the growth morphogen, and *b*_*u*_ is the cell growth constant.

The expression of the growth morphogen *m*_*G*_ is inhibited by a border morphogen *m*_*B*_, and both are subjected to advection, diffusion, production, and decay. As in (Ko & Lobo, 2019), we use the advective term to model how moving cells carry morphogens through space and the diffusive term to model how morphogens move within a tissue, such that

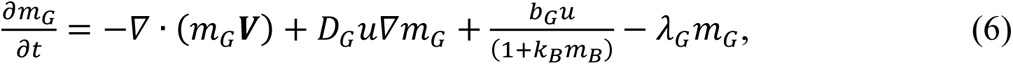

where *D*_*G*_ is the diffusion constant, *b*_*G*_ is the production constant, *k*_*B*_ is the inhibition constant of the border morphogen, and *λ*_*G*_ is the decay constant.

The border morphogen *m*_*B*_ is produced by cells at the border but inhibited by the anterior and posterior pole morphogens *m*_*A*_ and *m*_*P*_, respectively. When the concentration of *m*_*A*_ and *m*_*P*_ is zero at a given location, then the production rate constant of border morphogen is at its basal level *b*_*B*_. At higher concentrations of pole signals, the expression of the border morphogen is modulated according to the coefficient *α* that at negative values causes the pole signals to inhibit the expression of the border morphogen and, conversely, at positive values causes the pole signals to enhance the expression of the border morphogen (see Fig. 6B), as given by

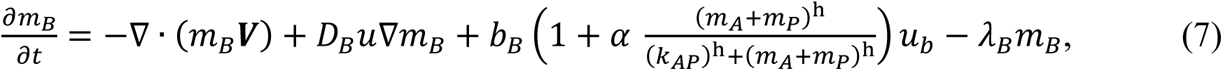

where *D*_*B*_ is the diffusion constant, *b*_*B*_ is the production constant, *k*_*AP*_ is the half-maximal concentration of the pole morphogens, and *λ*_*B*_ is the decay constant. The cells at the border are defined as *u*_*b*_ and computed by applying a Sobel edge detector to the cell density *u*.

The pole morphogens *m*_*A*_ and *m*_*P*_ are produced by the cells at the anterior pole *u*_*A*_ and the cells at the posterior pole *u*_*P*_, respectively, taking the form

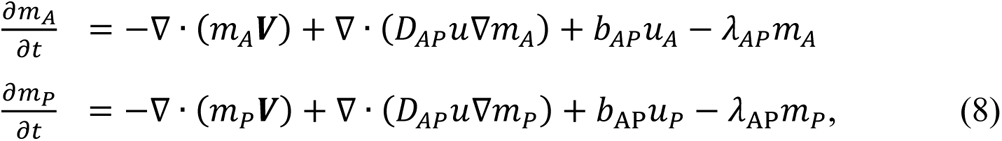

where *D*_*AP*_ is the diffusion constant, *b*_*AP*_ is the production constant, and *λ*_*AP*_ is the decay constant. The cells at the anterior and posterior poles, *u*_*A*_ and *u*_*P*_ respectively, are computed as the intersection of the border cells *u*_*B*_ and the vertical centerline of the top and bottom half of the domain, respectively (which always coincide with the centerline of the simulated worm shape due to symmetry).

### 4.4. Numerical methods

Simulations were computed in a two-dimensional domain in a uniform square lattice of 130 x 130 elements corresponding to 6.5 x 6.5 mm using the explicit upwind finite volume method with flux limiting and a zero flux boundary condition. The nonlocal integral term for adhesion is discretized into angular and radial components using bilinear interpolation, as in (Ko & Lobo, 2019). The system was numerically solved using ROWMAP (Weiner et al., 1997) and implemented in MATLAB (The MathWorks, Inc.). Body proportions of simulations were computed as the bounding box of cell density, where the cell density in border elements signifies sub-spatial-resolution width. The colors in the simulation panels and videos represent each variable normalized separately relative to its maximum value throughout space and time.

### 4.5. Machine learning method

We developed a novel machine learning approach to infer the model parameter values that can recapitulate the dynamics of experimental whole-body shapes by using a continuous descriptive model as an intermediary. The method is based on evolutionary computation, an optimization algorithm that mimics biological evolution. A population of 36 individuals each representing a different set of model parameters evolve by applying mutation, crossover, and selection operators towards a parameter set that can precisely recapitulate the shape dynamics in the descriptive model, and hence the experimental data.

The initial population is made of random individuals representing parameter sets from uniform distributions within the ranges defined in Supplementary Table 2. For each generation, 36 children are generated by randomly crossing all current individuals so that each pair of parents produces a pair of children with the same parameter sets, except each parameter can be swapped between the two children or replaced by a random one (using the same distribution), each event happening with a probably of 10%. All children (and parents in the first generation) are simulated with the mechanistic model using the parameter set they define, and a fitness is computed using the shape error between the simulation and the descriptive model. The new generation is selected as the 36 individuals among parents and children that have the best fitness (lowest error). The evolutionary algorithm runs for a predetermined time period, after which the best parameter set is returned.

The fitness of an individual simulation is computed using the dynamic shapes defined in the descriptive model. A shape error is defined as the average difference between each element in the simulated and descriptive shapes at a particular time point, expressed as

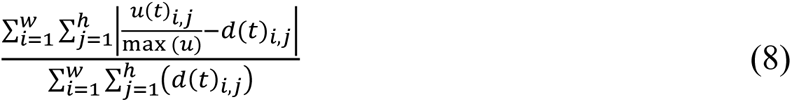

where *u*(*t*)_*i,j*_ is the simulated cell density at time *t* and location (*i*, *j*), and *d*(*t*)_*i,j*_ is the descriptive shape value at the same time and location. Notice that *u* ∈ [0, *k*_*c*_] and *d* ∈ [0,1].

To improve the performance of the evolutionary algorithm, the fitness of an individual is defined as the normalized simulated time before the shape error is higher than a threshold. In this way, simulations with high errors can be terminated earlier, resulting in a fitness less than 1, and only those staying within shape errors below the threshold are simulated for all the experimental data, resulting in a fitness of 1. Crucially, the shape error threshold changes dynamically during evolution. It starts with a high value of 1 but decreases when a new individual completes the whole simulation period (fitness of 1). The threshold is then updated to the maximum shape error during the simulation of this individual. In this way, the shape error threshold dynamically decreases as better models evolve and can complete the simulation. This increases the stringency of the fitness computation and hence reduces the computational cost of simulating models while increasing the evolutionary pressure towards better models. Additionally, simulations that have numerical errors or hit the boundary of the domain are also terminated early.

## Acknowledgments

We thank the members of the Lobo Lab and the planarian community for helpful discussions. This work was supported by the National Institute of General Medical Sciences of the National Institutes of Health under award number R35GM137953. The content is solely the responsibility of the authors and does not necessarily represent the official views of the National Institutes of Health. Computations used the UMBC High Performance Computing Facility (HPCF) supported by the NSF MRI program grants OAC-1726023, CNS-0821258, and CNS-1228778, and the SCREMS program grant DMS-0821311.

## Author contributions

J.M.K. and D.L. designed the study, developed the models, analyzed the data, and wrote the manuscript; W.R. performed the *in vivo* experiments; J.M.K. performed the simulations; D.L. secured funding.

## Data availability

The source code is freely available from GitHub (https://github.com/lobolab/planarian-growth-degrowth).

## References

Adell, T., Saló, E., Boutros, M., & Bartscherer, K. (2009). Smed-Evi/Wntless is required for beta-catenin-dependent and -independent processes during planarian regeneration. Development, 136, 905–910. 10.1242/dev.033761

Almuedo-Castillo, M., Crespo, X., Seebeck, F., Bartscherer, K., Salò, E., & Adell, T. (2014). JNK Controls the Onset of Mitosis in Planarian Stem Cells and Triggers Apoptotic Cell Death Required for Regeneration and Remodeling. PLOS Genetics, 10(6), e1004400. 10.1371/journal.pgen.1004400

Baguñà, J., Romero, R., Saló, E., Collet, J., Auladell, C., Ribas, M., Riutort, M., García-Fernàndez, J., Burgaya, F., & Bueno, D. (1990). Growth, Degrowth and Regeneration as Developmental Phenomena in Adult Freshwater Planarians. In H.-J. Marthy (Ed.), Experimental Embryology in Aquatic Plants and Animals (pp. 129–162). Springer US. 10.1007/978-1-4615-3830-1_7

Birkholz, T. R., Van Huizen, A. V., & Beane, W. S. (2019). Staying in shape: Planarians as a model for understanding regenerative morphology. Seminars in Cell and Developmental Biology, 87, 105–115. 10.1016/j.semcdb.2018.04.014

Bonar, N. A., Gittin, D. I., & Petersen, C. P. (2022). Src acts with WNT/FGFRL signaling to pattern the planarian anteroposterior axis. Development, 149(7), dev200125. 10.1242/dev.200125

Cebrià, F., Guo, T., Jopek, J., & Newmark, P. A. (2007). Regeneration and maintenance of the planarian midline is regulated by a slit orthologue. Developmental Biology, 307(2), 394–406. 10.1016/j.ydbio.2007.05.006

Clark, E. G., & Petersen, C. P. (2023). BMP suppresses WNT to integrate patterning of orthogonal body axes in adult planarians (p. 2023.01.10.523528). bioRxiv. 10.1101/2023.01.10.523528

Crocker, J., Ilsley, G. R., & Stern, D. L. (2016). Quantitatively predictable control of Drosophila transcriptional enhancers in vivo with engineered transcription factors. Nature Genetics, 48(3), 292–298. 10.1038/ng.3509

Dillon, R., Gadgil, C., & Othmer, H. G. (2003). Short- and long-range effects of Sonic hedgehog in limb development. Proceedings of the National Academy of Sciences, 100(18), 10152– 10157. 10.1073/pnas.1830500100

Felix, D. A., Gutiérrez-Gutiérrez, Ó., Espada, L., Thems, A., & González-Estévez, C. (2019). It is not all about regeneration: Planarians striking power to stand starvation. Seminars in Cell and Developmental Biology, 87, 169–181. 10.1016/j.semcdb.2018.04.010

Fraguas, S., Barberán, S., & Cebrià, F. (2011). EGFR signaling regulates cell proliferation, differentiation and morphogenesis during planarian regeneration and homeostasis. Developmental Biology, 354(1), 87–101. 10.1016/j.ydbio.2011.03.023

Francois, P., & Siggia, E. D. (2010). Predicting embryonic patterning using mutual entropy fitness and in silico evolution. Development, 137, 2385–2395. 10.1242/Dev.048033

Gaviño, M. A., & Reddien, P. W. (2011). A Bmp/Admp Regulatory Circuit Controls Maintenance and Regeneration of Dorsal-Ventral Polarity in Planarians. Current Biology, 21(4), 294–299. 10.1016/j.cub.2011.01.017

Germann, P., Marin-Riera, M., & Sharpe, J. (2019). ya||a: GPU-Powered Spheroid Models for Mesenchyme and Epithelium. Cell Systems, 8(3), 261–266.e3. 10.1016/j.cels.2019.02.007

González-Estévez, C., Felix, D. A., Rodríguez-Esteban, G., & Aboobaker, A. A. (2012a). Decreased neoblast progeny and increased cell death during starvation-induced planarian degrowth. International Journal of Developmental Biology, 56(1-2–3), Article 1-2–3. 10.1387/ijdb.113452cg

González-Estévez, C., Felix, D. A., Rodríguez-Esteban, G., & Aboobaker, A. A. (2012b). Decreased neoblast progeny and increased cell death during starvation-induced planarian degrowth. The International Journal of Developmental Biology, 56, 83–91. 10.1387/ijdb.113452cg

González-Estévez, C., Felix, D. A., Smith, M. D., Paps, J., Morley, S. J., James, V., Sharp, T. V., & Aboobaker, A. A. (2012). SMG-1 and mTORC1 Act Antagonistically to Regulate Response to Injury and Growth in Planarians. PLoS Genetics, 8, e1002619. 10.1371/journal.pgen.1002619

Gurley, K. A., Elliott, S. A., Simakov, O., Schmidt, H. A., Holstein, T. W., & Sánchez Alvarado, A. (2010). Expression of secreted Wnt pathway components reveals unexpected complexity of the planarian amputation response. Developmental Biology, 347(1), 24–39. 10.1016/j.ydbio.2010.08.007

Gurley, K. A., Rink, J. C., & Alvarado, A. S. (2008). β-Catenin Defines Head Versus Tail Identity During Planarian Regeneration and Homeostasis. Science, 319(5861), 323–327. 10.1126/science.1150029

Harmansa, S., & Lecuit, T. (2021). Forward and feedback control mechanisms of developmental tissue growth. Cells & Development, 168, 203750. 10.1016/j.cdev.2021.203750

Herath, S., & Lobo, D. (2020). Cross-inhibition of Turing patterns explains the self-organized regulatory mechanism of planarian fission. Journal of Theoretical Biology, 485, 110042. 10.1016/j.jtbi.2019.110042

Hill, E. M., & Petersen, C. P. (2015). Wnt/Notum spatial feedback inhibition controls neoblast differentiation to regulate reversible growth of the planarian brain. Development, 142(24), 4217–4229. 10.1242/dev.123612

Kicheva, A., & Briscoe, J. (2023). Control of Tissue Development by Morphogens. Annual Review of Cell and Developmental Biology, 39(1), null. 10.1146/annurev-cellbio-020823-011522

Ko, J. M., & Lobo, D. (2019). Continuous Dynamic Modeling of Regulated Cell Adhesion: Sorting, Intercalation, and Involution. Biophysical Journal, 2166–2179. 10.1016/j.bpj.2019.10.032

Ko, J. M., Mousavi, R., & Lobo, D. (2022). Computational Systems Biology of Morphogenesis. In S. Cortassa & M. A. Aon (Eds.), Computational Systems Biology in Medicine and Biotechnology: Methods and Protocols (pp. 343–365). Springer US. 10.1007/978-1-0716-1831-8_14

Lapan, S. W., & Reddien, P. W. (2011). Dlx and sp6-9 Control Optic Cup Regeneration in a Prototypic Eye. PLOS Genetics, 7(8), e1002226. 10.1371/journal.pgen.1002226

Lobo, D., Beane, W. S., & Levin, M. (2012). Modeling planarian regeneration: A primer for reverse-engineering the worm. PLoS Computational Biology, 8, e1002481. 10.1371/journal.pcbi.1002481

Lobo, D., & Levin, M. (2015). Inferring Regulatory Networks from Experimental Morphological Phenotypes: A Computational Method Reverse-Engineers Planarian Regeneration. PLOS Computational Biology, 11(6), e1004295. 10.1371/journal.pcbi.1004295

Lobo, D., & Levin, M. (2017). Computing a Worm: Reverse-Engineering Planarian Regeneration. In A. Adamatzky (Ed.), Advances in Unconventional Computing. Volume 2: Prototypes, Models and Algorithms (pp. 637–654). Springer International Publishing. 10.1007/978-3-319-33921-4_24

Lobo, D., Malone, T. J., & Levin, M. (2013). Planform: An application and database of graph-encoded planarian regenerative experiments. Bioinformatics, 29, 1098–1100. 10.1093/bioinformatics/btt088

Lobo, D., Morokuma, J., & Levin, M. (2016). Computational discovery and in vivo validation of hnf4 as a regulatory gene in planarian regeneration. Bioinformatics, 32(17), 2681–2685. 10.1093/bioinformatics/btw299

Marin-Riera, M., Brun-Usan, M., Zimm, R., V??likangas, T., & Salazar-Ciudad, I. (2015). Computational modeling of development by epithelia, mesenchyme and their interactions: A unified model. Bioinformatics, 32(2), 219–225. 10.1093/bioinformatics/btv527

Meinhardt, H. (1982). Models of Biological Pattern Formation. Academic Press.

Meinhardt, H. (2004). Different strategies for midline formation in bilaterians. Nature Reviews Neuroscience, 5(6), 502–510. 10.1038/nrn1410

Merryman, M. S., Alvarado, A. S., & Jenkin, J. C. (2018). Culturing Planarians in the Laboratory. In J. C. Rink (Ed.), Planarian Regeneration: Methods and Protocols (pp. 241–258). Springer. 10.1007/978-1-4939-7802-1_5

Miikkulainen, R., & Forrest, S. (2021). A biological perspective on evolutionary computation. Nature Machine Intelligence, 3(1), 9–15. 10.1038/s42256-020-00278-8

Miller, C. M., & Newmark, P. A. (2012). An insulin-like peptide regulates size and adult stem cells in planarians. The International Journal of Developmental Biology, 56, 75–82. 10.1387/ijdb.113443cm

Mirams, G. R., Arthurs, C. J., Bernabeu, M. O., Bordas, R., Cooper, J., Corrias, A., Davit, Y., Dunn, S.-J., Fletcher, A. G., Harvey, D. G., Marsh, M. E., Osborne, J. M., Pathmanathan, P., Pitt-Francis, J., Southern, J., Zemzemi, N., & Gavaghan, D. J. (2013). Chaste: An Open Source C++ Library for Computational Physiology and Biology. PLoS Computational Biology, 9(3), e1002970. 10.1371/journal.pcbi.1002970

Molina, M. D., Saló, E., & Cebrià, F. (2007). The BMP pathway is essential for re-specification and maintenance of the dorsoventral axis in regenerating and intact planarians. Developmental Biology, 311(1), 79–94. 10.1016/j.ydbio.2007.08.019

Mousavi, R., Konuru, S. H., & Lobo, D. (2021). Inference of dynamic spatial GRN models with multi-GPU evolutionary computation. Briefings in Bioinformatics, 22(5), 1–11. 10.1093/bib/bbab104

Oviedo, N. J., Newmark, P. A., & Alvarado, A. S. (2003). Allometric scaling and proportion regulation in the freshwater planarian Schmidtea mediterranea. Developmental Dynamics, 226(2), 326–333. 10.1002/dvdy.10228

Pascual-Carreras, E., Herrera-Úbeda, C., Rosselló, M., Coronel-Córdoba, P., Garcia-Fernàndez, J., Saló, E., & Adell, T. (2021). Analysis of Fox genes in Schmidtea mediterranea reveals new families and a conserved role of Smed-foxO in controlling cell death. Scientific Reports, 11(1), Article 1. 10.1038/s41598-020-80627-0

Pascual-Carreras, E., Marin-Barba, M., Herrera-Úbeda, C., Font-Martín, D., Eckelt, K., de Sousa, N., García-Fernández, J., Saló, E., & Adell, T. (2020). Planarian cell number depends on blitzschnell, a novel gene family that balances cell proliferation and cell death. Development, 147(7), dev184044. 10.1242/dev.184044

Peiris, T. H., Weckerle, F., Ozamoto, E., Ramirez, D., Davidian, D., Garcia-Ojeda, M. E., & Oviedo, N. J. (2012). TOR signaling regulates planarian stem cells and controls localized and organismal growth. Journal of Cell Science, 125, 1657–1665. 10.1242/jcs.104711

Pellettieri, J., Fitzgerald, P., Watanabe, S., Mancuso, J., Green, D. R., & Sánchez Alvarado, A. (2010). Cell death and tissue remodeling in planarian regeneration. Developmental Biology, 338(1), 76–85. 10.1016/j.ydbio.2009.09.015

Petersen, C. P., & Reddien, P. W. (2009). A wound-induced Wnt expression program controls planarian regeneration polarity. Proceedings of the National Academy of Sciences, 106(40), 17061–17066. 10.1073/pnas.0906823106

Petersen, C. P., & Reddien, P. W. (2011). Polarized notum Activation at Wounds Inhibits Wnt Function to Promote Planarian Head Regeneration. Science, 332(6031), 852–855. 10.1126/science.1202143

Pietak, A., Bischof, J., LaPalme, J., Morokuma, J., & Levin, M. (2019). Neural control of body-plan axis in regenerating planaria. PLOS Computational Biology, 15(4), e1006904. 10.1371/journal.pcbi.1006904

Reddien, P. W. (2021). Positional Information and Stem Cells Combine to Result in Planarian Regeneration. Cold Spring Harbor Perspectives in Biology, a040717. 10.1101/cshperspect.a040717

Reddien, P. W., Bermange, A. L., Kicza, A. M., & Sanchez Alvarado, A. (2007). BMP signaling regulates the dorsal planarian midline and is needed for asymmetric regeneration. Development, 134(22), 4043–4051. 10.1242/dev.007138

Schad, E. G., & Petersen, C. P. (2020). STRIPAK Limits Stem Cell Differentiation of a WNT Signaling Center to Control Planarian Axis Scaling. Current Biology, 30(2), 254–263.e2. 10.1016/j.cub.2019.11.068

Schiffmann, Y. (2011). Turing-Child field underlies spatial periodicity in Drosophila and planarians. Progress in Biophysics and Molecular Biology, 105, 258–269.

Schnell, S., Maini, P., Newman, S. A., & Newman, T. (2007). Multiscale Modeling of Developmental Systems. In G. Schatten (Ed.), Current Topics in Developmental Biology (Vol. 81). Elsevier.

Scimone, M. L., Cote, L. E., Rogers, T., & Reddien, P. W. (2016). Two FGFRL-Wnt circuits organize the planarian anteroposterior axis. eLife, 5, e12845. 10.7554/eLife.12845

Scimone, M. L., Kravarik, K. M., Lapan, S. W., & Reddien, P. W. (2014). Neoblast Specialization in Regeneration of the Planarian Schmidtea mediterranea. Stem Cell Reports, 3(2), 339–352. 10.1016/j.stemcr.2014.06.001

Sharpe, J. (2017). Computer modeling in developmental biology: Growing today, essential tomorrow. Development, 144(23), 4214–4225. 10.1242/dev.151274

Stückemann, T., Cleland, J. P., Werner, S., Thi-Kim Vu, H., Bayersdorf, R., Liu, S.-Y., Friedrich, B., Jülicher, F., & Rink, J. C. (2017). Antagonistic Self-Organizing Patterning Systems Control Maintenance and Regeneration of the Anteroposterior Axis in Planarians. Developmental Cell, 40(3), 248–263.e4. 10.1016/j.devcel.2016.12.024

Sureda-Gomez, M., & Adell, T. (2019). Planarian organizers. Seminars in Cell & Developmental Biology, 87, 95–104. 10.1016/j.semcdb.2018.05.021

Sureda-Gómez, M., Martín-Durán, J. M., & Adell, T. (2016). Localization of planarian β-CATENIN-1 reveals multiple roles during anterior-posterior regeneration and organogenesis. Development, 143(22), 4149–4160. 10.1242/dev.135152

Takeda, H., Nishimura, K., & Agata, K. (2009). Planarians maintain a constant ratio of different cell types during changes in body size by using the stem cell system. Zoological Science, 26, 805–813.

Texada, M. J., Koyama, T., & Rewitz, K. (2020). Regulation of body size and growth control. Genetics, 216(2), 269–313. 10.1534/genetics.120.303095

Thommen, A., Werner, S., Frank, O., Philipp, J., Knittelfelder, O., Quek, Y., Fahmy, K., Shevchenko, A., Friedrich, B. M., Jülicher, F., & Rink, J. C. (2019). Body size-dependent energy storage causes Kleiber’s law scaling of the metabolic rate in planarians. eLife, 8, e38187. 10.7554/eLife.38187

Uzkudun, M., Marcon, L., & Sharpe, J. (2015). Data-driven modelling of a gene regulatory network for cell fate decisions in the growing limb bud. Molecular Systems Biology, 11(7), 815. 10.15252/msb.20145882

Weiner, R., Schmitt, B. A., & Podhaisky, H. (1997). ROWMAP—a ROW-code with Krylov techniques for large stiff ODEs. Applied Numerical Mathematics, 25(2–3), 303–319.

Werner, S., Stückemann, T., Beirán Amigo, M., Rink, J. C., Jülicher, F., Friedrich, B. M., Amigo, M. B., Rink, J. C., Jülicher, F., & Friedrich, B. M. (2015). Scaling and regeneration of self-organized patterns. Physical Review Letters, 114(13), 138101. 10.1103/PhysRevLett.114.138101

Witchley, J. N., Mayer, M., Wagner, D. E., Owen, J. H., & Reddien, P. W. (2013). Muscle Cells Provide Instructions for Planarian Regeneration. Cell Reports, 4(4), 633–641. 10.1016/j.celrep.2013.07.022

Ziman, B., Karabinis, P., Barghouth, P., & Oviedo, N. J. (2020). Sirtuin-1 regulates organismal growth by altering feeding behavior and intestinal morphology in planarians. Journal of Cell Science, 133(10), jcs239467. 10.1242/jcs.239467

