## Supplementary Information for "Mechanistic Regulation of Planarian Shape During Growth and Degrowth"

1. Supplementary tables ... 2
2. Supplementary figures ... 4
3. Supplementary videos ... 6

### 1. Supplementary tables

**Supplementary Table 1.** Parameters used for testing the mechanistic model (Fig. 3).

| Param. | Description | Value | Units | Source |
|---|---|---|---|---|
| $D_{AP}$ | Pole morphogens diffusion constant | 50 | $\mu m^2 \cdot s^{-1}$ | Estimated |
| $b_{AP}$ | Pole morphogens production constant | 1000 | $amol \cdot cell^{-1} \cdot h^{-1}$ | Estimated |
| $\lambda_{AP}$ | Pole morphogens decay constant | 0.1 | $h^{-1}$ | Estimated |
| $k_{AP}$ | Poles half-maximal concentration constant | 0.5 | $amol \cdot \mu m^{-2}$ | Estimated |
| $\alpha$ | Poles coefficient | -1 | dimensionless | Estimated |
| $D_B$ | Border morphogen diffusion constant | 50 | $\mu m^2 \cdot s^{-1}$ | Estimated |
| $b_B$ | Border morphogen production constant | 80 | $amol \cdot cell^{-1} \cdot h^{-1}$ | Estimated |
| $\lambda_B$ | Border morphogen decay constant | 0.08 | $h^{-1}$ | Estimated |
| $k_B$ | Border morphogen inhibition constant | 1 | $\mu m^2 \cdot amol^{-1}$ | Estimated |
| $D_G$ | Growth morphogen diffusion constant | 0 | $\mu m^2 \cdot s^{-1}$ | Estimated |
| $b_G$ | Growth morphogen production constant | 50 | $amol \cdot cell^{-1} \cdot h^{-1}$ | Estimated |
| $\lambda_G$ | Growth morphogen decay constant | 0.2 | $h^{-1}$ | Estimated |
| $k_G$ | Growth morphogen half-maximal concentration constant | 0.5 | $amol \cdot \mu m^{-2}$ | Estimated |
| $\lambda$ | Apoptosis rate constant | 0.01 | $h^{-1}$ | Estimated |
| $k_p$ | Cell dispersion constant | 35 | $\frac{\mu m^2 \cdot s^{-1}}{cell \cdot \mu m^{-2}}$ | Estimated |
| $k_a$ | Cell adhesion constant | 15 | $\frac{\mu m^2 \cdot s^{-1}}{cell \cdot \mu m^{-2}}$ | Estimated |
| $R$ | Radius of adhesion forces | 100 | $\mu m$ | HEK293 cells (Ko & Lobo, 2019) |
| $\Phi$ | Constant of proportionality | 100 | dimensionless | Estimated |
| $b_u$ | Tissue growth rate constant | 1/12 | $h^{-1}$ | HEK293 cells (Ko & Lobo, 2019) |
| $k_c$ | Cell carrying capacity | 0.00559504 | $cell \cdot \mu m^{-2}$ | HEK293 cells (Ko & Lobo, 2019) |
| $u_0$ | Initial cell density | $k_c$ | $cell \cdot \mu m^{-2}$ | (Ko & Lobo, 2019) |

**Supplementary Table 2.** Parameters inferred and ranges used with the evolutionary algorithm (Fig. 5). Rest of model parameters as in Supplementary Table 1.

| **Param.** | **Description** | **Value** | **Min** | **Max** | **Units** |
|---|---|---|---|---|---|
| $D_{AP}$ | Pole morphogens diffusion constant | 29.98 | 1.66 | 30 | $\mu m^2 \cdot s^{-1}$ |
| $b_{AP}$ | Pole morphogens production constant | 2,511.05 | 0 | 10,000 | $amol \cdot cell^{-1} \cdot h^{-1}$ |
| $\lambda_{AP}$ | Pole morphogens decay constant | 0.037 | 0.0 | 1.0 | $h^{-1}$ |
| $k_{AP}$ | Poles half-maximal concentration constant | 0.24 | 0.1 | 1.0 | $amol \cdot \mu m^{-2}$ |
| $D_B$ | Border morphogen diffusion constant | 7.59 | 1.66 | 30 | $\mu m^2 \cdot s^{-1}$ |
| $b_B$ | Border morphogen production constant | 9,599.93 | 0 | 10,000 | $amol \cdot cell^{-1} \cdot h^{-1}$ |
| $\lambda_B$ | Border morphogen decay constant | 0.040 | 0.0 | 1.0 | $h^{-1}$ |
| $D_G$ | Growth morphogen diffusion constant | 25.40 | 1.66 | 30 | $\mu m^2 \cdot s^{-1}$ |
| $b_G$ | Growth morphogen production constant | 8,917.35 | 0 | 10,000 | $amol \cdot cell^{-1} \cdot h^{-1}$ |
| $\lambda_G$ | Growth morphogen decay constant | 0.27 | 0.0 | 1.0 | $h^{-1}$ |
| $k_G$ | Growth morphogen half-maximal concentration constant | 0.20 | 0.0 | 1.0 | $amol \cdot \mu m^{-2}$ |
| $\lambda$ | Apoptosis rate constant | 0.036 | 0.0 | 0.1 | $h^{-1}$ |
| $k_p$ | Cell dispersion constant | 26.73 | 0 | 70 | $\frac{\mu m^2 \cdot s^{-1}}{cell \cdot \mu m^{-2}}$ |
| $k_a$ | Cell adhesion constant | 27.32 | 5 | 120 | $\frac{\mu m^2 \cdot s^{-1}}{cell \cdot \mu m^{-2}}$ |
| $u_0$ | Initial cell density | $0.41 \cdot k_c$ | $0.0 \cdot k_c$ | $1.0 \cdot k_c$ | $cell \cdot \mu m^{-2}$ |

### 2. Supplementary figures

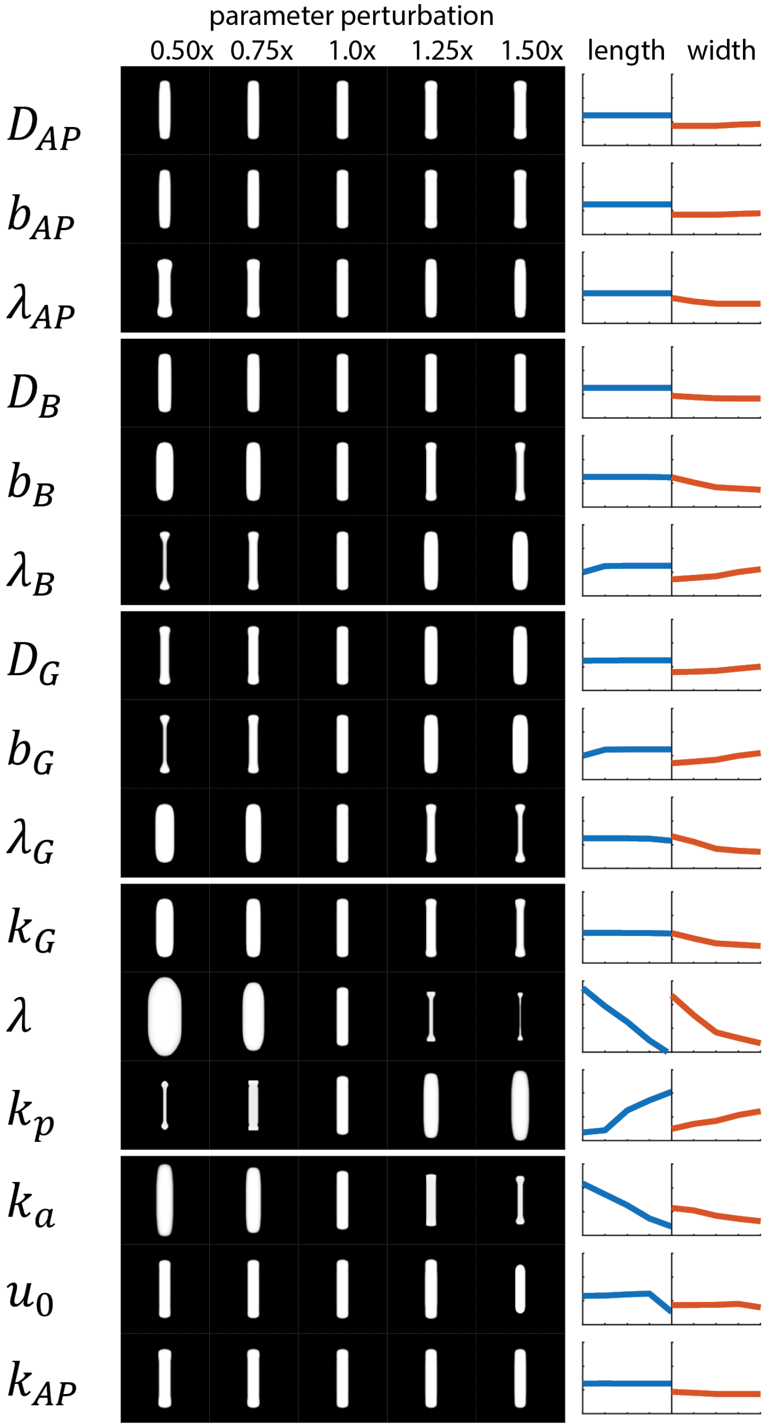


**Supplementary Figure 1.** Sensitivity analysis of the mechanistic model with the evolved parameters for planarian growth dynamics. Each parameter was perturbed by multiplying its value by a specific factor. The resultant whole-body shape is shown after three simulated weeks. Line plots show the length (blue) and width (orange) of the whole-body shapes in each row (length plot range: 3 to 6 mm; width plot range: 0 to 3 mm).

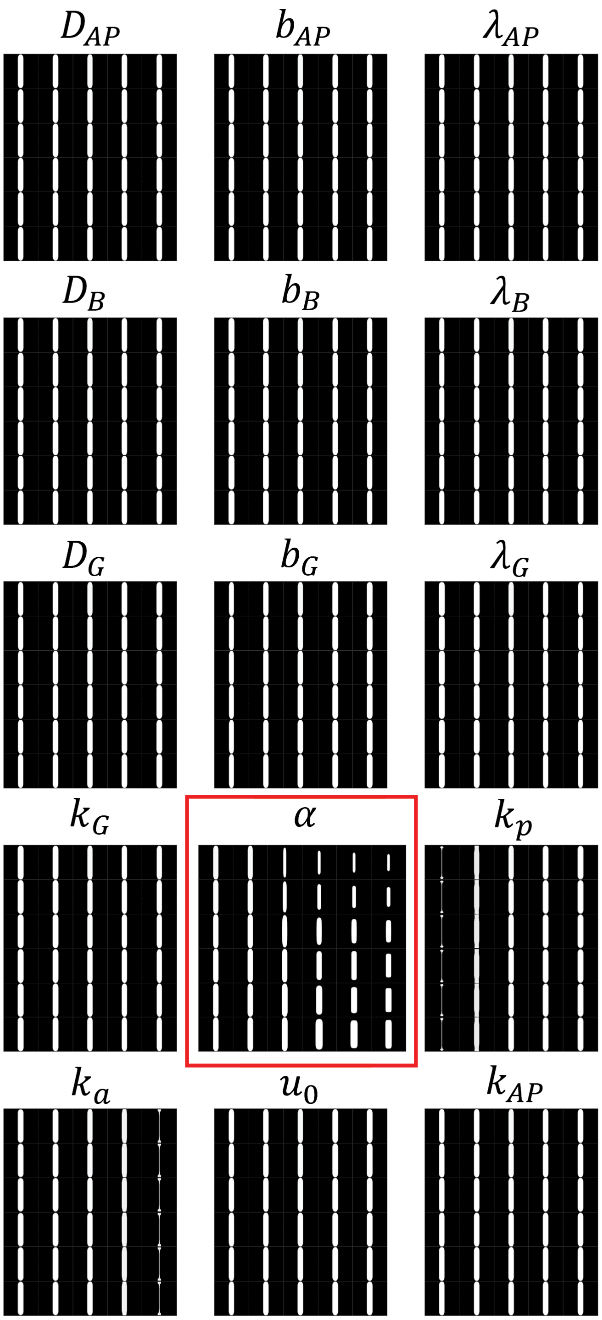


**Supplementary Figure 2.** Simulations varying the apoptosis rate constant and one other parameter reveal that only the combination of apoptosis rate and pole coefficient can induce degrowth behaviors. Each panel shows final cell density for degrowth simulations (Fig. 7) using the inferred parameters for growth except cell apoptosis ($\lambda$) and one other parameter. The value of $\lambda$ varies vertically from 1.0x (bottom) to 1.1x (top) in increments of 0.02x. The value of the parameter indicated in the panels varies horizontally as in Supplementary Figure 1, except the pole coefficient $\alpha$ (red box), which varies from -1 to 1 in increments of 0.4. Simulations are terminated earlier when cell density reaches the boundary of the simulation domain, indicating growth instead of degrowth.

### 3. Supplementary videos

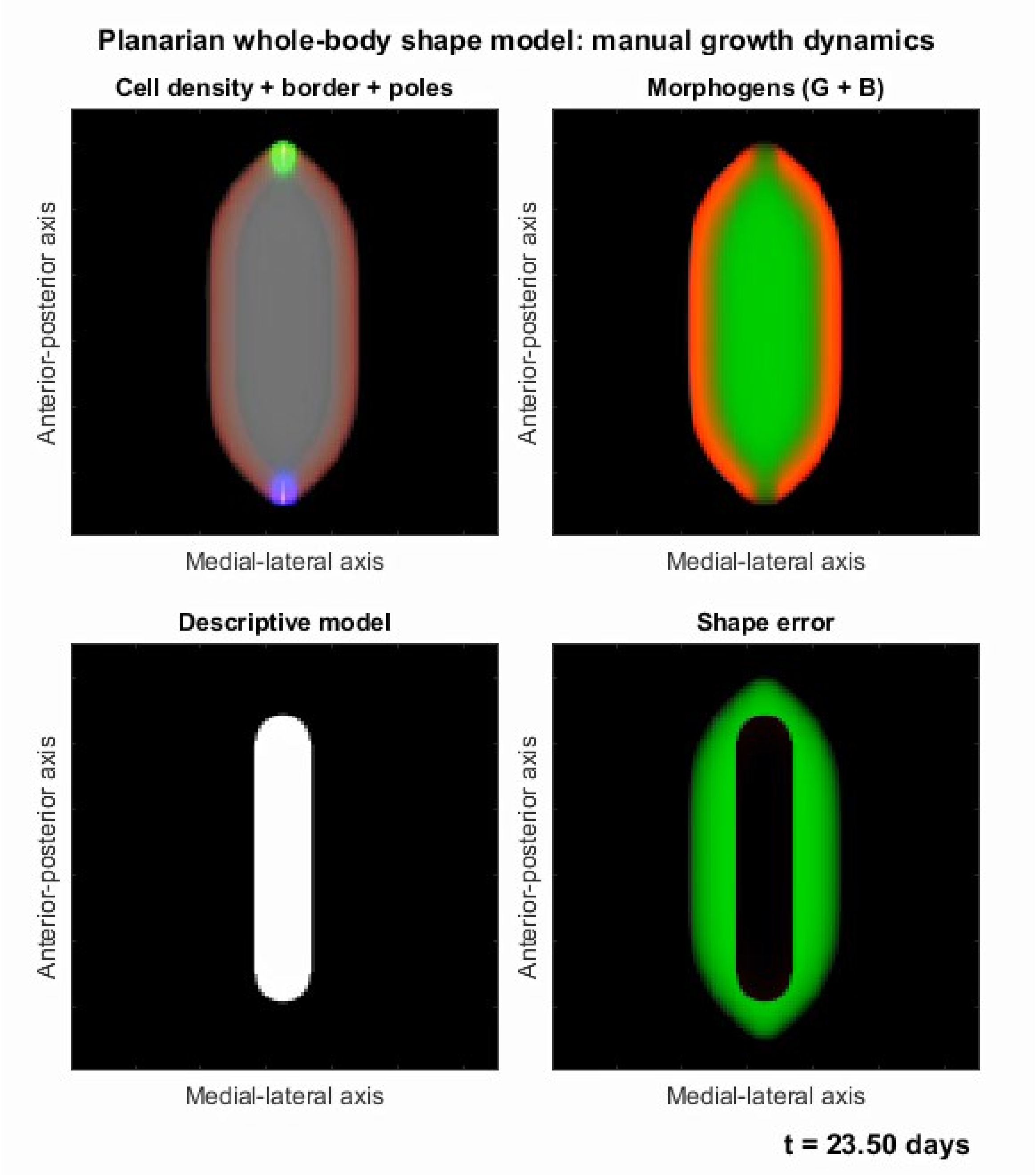


**Supplementary Video 1.** Simulation comparing the mechanistic model of planarian whole-body shape using a manually estimated parameter set with the descriptive model of growth derived from experimental data, as shown in Figure 3.

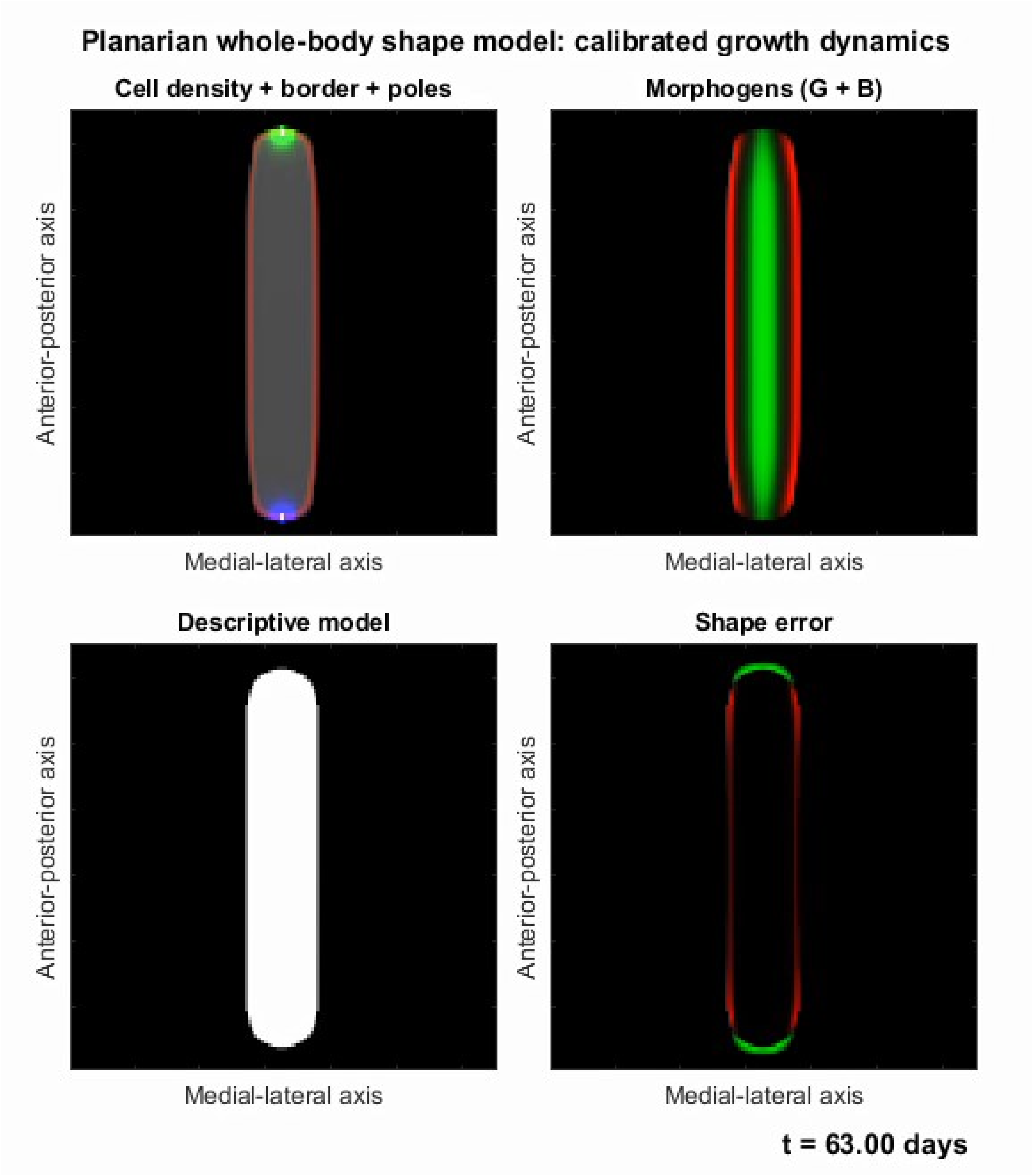


**Supplementary Video 2.** Simulation comparing the mechanistic model of planarian whole-body shape using the parameter set calibrated by the machine learning with the descriptive model of growth derived from experimental data, as shown in Figure 5.

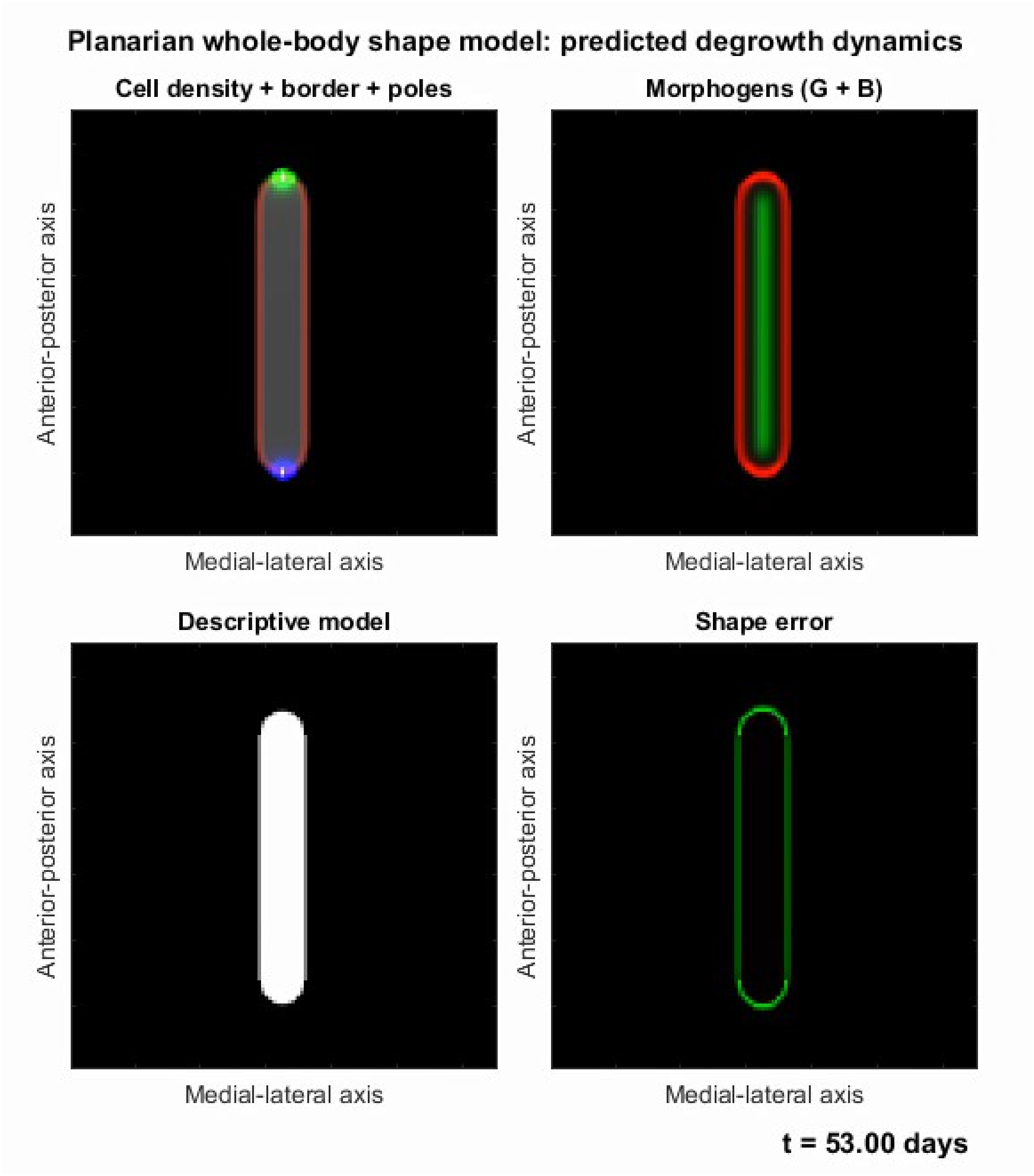


**Supplementary Video 3.** Simulation comparing the calibrated mechanistic model of planarian whole-body shape with higher apoptosis rate and pole coefficient with the descriptive model of degrowth derived from experimental data, as shown in Figure 7.
